# BRCA2 and RAD54B FxPP motifs Bind DMC1 Filaments through Persistent and Transient Interfaces

**DOI:** 10.64898/2026.07.24.740592

**Authors:** Pauline Dupaigne, Simona Miron, Sonia Baconnais, Pierre Legrand, Sari E. van Rossum-Fikkert, Koichi Sato, Atifa Majeed, Marine Le Hingrat, Malika Ouldali, Alex N. Zelensky, Roland Kanaar, Philippe Cuniasse, Sophie Zinn-Justin

## Abstract

During meiosis, double-strand DNA breaks are programmed to initiate homologous recombination. The DMC1 recombinogenic filament is central to the repair of these breaks. However, the three-dimensional structures of the protein-protein interfaces regulating the assembly and activity of this filament have not been described yet. We and others previously reported that in BRCA2, a P-motif called PhePP binds to DMC1 in its oligomeric and filament states. Here we identified a similar P-motif in the DNA translocase RAD54B. We solved the cryo-electron microscopy (cryo-EM) structures of BRCA2 and RAD54B P-motif peptides bound to a ssDNA-DMC1 filament at 1.9-2.0 Å resolution. While these peptides only share the sequence F-[IV]-P-P, they bind to the filament through larger 9-10 residue core sequences with superimposable structures. Both peptides bind to a hydrophobic and negatively charged site, named the P-site, on a single DMC1 protomer and stabilize the ssDNA-DMC1 filament. Mutagenesis experiments and molecular dynamics simulations identified additional transient interactions between positively charged residues of the peptides and negatively charged patches distributed on the DMC1 protomers. We propose that BRCA2 and RAD54B stably dock at the P-site of a single DMC1 protomer but also bridge two DMC1 protomers within the presynaptic filament via transient contacts with the adjacent protomer.

## Introduction

During meiosis, haploid gametes are generated with new combinations of genetic materials. Homologous recombination (HR) contributes to genetic variation by shuffling preexisting sequences. It drives chromosome crossovers, ensuring proper homolog segregation, generation of genetic diversity and faithful transmission of genetic information (1). Failure in meiotic recombination can result in chromosome mis-segregation, aneuploidy and defective gamete formation, leading to infertility and genomic disorders (2). The central step of HR is the homology search, wherein RecA/RAD51 strand exchange proteins, also called recombinases, replace single-stranded DNA (ssDNA) binding proteins on ssDNA and catalyze the invasion of a double-stranded DNA (dsDNA) to find and align homologous sequences (3).

Eukaryotic recombinases exhibit a two-lobed architecture with a disordered N-terminus, an α-helical N-terminal domain of about 60 residues, a linker of about 15 residues containing a single α-helix and a large C-terminal ATPase domain of about 240 residues with DNA interacting loops (4–6). Structural analyses of human RAD51 nucleoprotein filaments revealed that the RAD51 ATPase domain establishes extensive contacts between neighboring subunits (7–9). Upon filament assembly, two adjacent ATPase domains form an ATP binding cavity in which one protomer directly binds to the phosphates of ATP while the other protomer interacts with its adenosine moiety (9). In addition, the linker region of one RAD51 protomer inserts into a hydrophobic pocket of the ATPase domain of the adjacent protomer. This interaction is essential as mutating the conserved F86 in the linker into glutamic acid prevents RAD51 self-association and foci formation (4). Within the filaments, the DNA is stretched and stabilized, making it more conducive to homology search (8, 9). The N-terminal region of RAD51 facilitates the formation of the nucleoprotein filament and is essential for strand exchange (10, 11). Within the filament, it interacts with both the ATPase domain of its own protomer and that of the adjacent protomer. During the homology search, it directly binds to the dsDNA to assemble the D-loop, essential for the processing of the double-strand break (DSB) (11, 12).

In meiosis, RAD51 acts together with an additional recombinase, DMC1, which exhibits unique properties: it preferentially promotes interhomolog recombination over sister-chromatid exchange, an aspect essential for accurate genetic distribution to gametes and proper meiotic chromosome segregation (13). RAD51 and DMC1 are both required for mammalian meiosis (14–16). In humans, they share 54 % of sequence identity; RAD51 forms heterogeneous oligomers (17), whereas DMC1 mainly assembles into octamers (18). Upon DNA binding, both recombinases produce similar nucleoprotein filaments (19, 20). Current evidence suggests that DMC1 is the primary strand-exchange enzyme during mammalian meiosis, whereas RAD51 binds further from the break and has an accessory (even if essential) function in meiotic recombination (21). Consistently, DMC1 contains specific residues in loop 2 that could be responsible for its mismatch tolerance (19, 20).

Both RAD51 and DMC1 are regulated by the central HR mediator BRCA2 (22). Indeed, absence of BRCA2 reduces the number of RAD51 and DMC1 punctate immunostaining signals (or foci) observed on meiotic chromosomes and results in complete infertility (23, 24). BRCA2 plays an essential role in favoring the loading of RAD51 and DMC1 on resected ssDNA at the DSBs. It can also stabilize RAD51 and DMC1 nucleoprotein filaments. At the molecular level, BRCA2 exhibits 8 BRC repeats (also named A-motifs) that bind to monomeric recombinases (4, 25, 26). Through these repeats, BRCA2 acts as an assembly chaperone that promotes nucleation of the recombinogenic filaments. In addition, BRCA2 possesses two identified conserved motifs sharing the sequence F-x-P-P (also named P-motifs) that interact with these filaments: the TR2 motif (encoded by exon 27) that binds to ssDNA- and dsDNA-RAD51 (10, 27) and the PhePP motif (28) (encoded by exon 14), located in the meiosis-specific region of BRCA2 (29), which binds to ssDNA- and dsDNA-DMC1 (30, 31).

Several structural studies (30–33) have now demonstrated that P-motifs represent a conserved mechanism for binding the oligomeric form of recombinases, which likely dates to the origins of eukaryotes, and that such binding can stabilize the recombinogenic filaments. The prototypical P-motifs were identified experimentally using *in vitro* interaction studies and mutagenesis. Extending the list of P-motif-containing proteins is complicated, because the pattern [FW]-x-P-P (defined from the BRCA2 and RAD51AP1 motifs) or [FW]-x-x-P (if the *S. cerevisiae* RAD54 P-motif is included) is very common. Therefore, novel P-motifs have mostly been identified in protein regions known to interact with a recombinase. For example, in our previous work (30) we spotted a conserved F-x-P-P consensus in the RAD51-binding domain (FRBD) of the FIGNL1 “antirecombinase” protein, where it is positioned next to the BRC-like A-motif, essential for RAD51 binding. We confirmed that the FRBD P-motif can weakly interact with DMC1. Similarly, motifs with F-x-x-P consensus have been previously spotted in Brh2 (fungal BRCA2 ortholog) (34, 35). Structural analyses of the P-motif complexes revealed that their interactions with recombinases extend beyond the [FW]-x-P-P sequence (30–32). Incorporating this information into bioinformatic analyses could allow unbiased detection of recombinase-binding proteins based on sequence alone.

The SWI2/SNF2-family DNA translocases RAD54 and RAD54B are important regulators of RAD51 and DMC1 during HR. These ATP-driven motor proteins stimulate recombinase-driven strand exchange (36–39). The helicase domains of human RAD54 and RAD54B share 48% of sequence identity. However, they possess mostly disordered divergent N-terminal regions that directly bind to RAD51 (36, 38, 40). The N-terminal domain of RAD54B, from S26 to R225, also directly binds to DMC1 (38). Mice lacking either RAD54, RAD54B or both RAD54 and RAD54B are fertile (39). Mice lacking RAD54 still exhibit meiotic abnormalities, including persistent RAD51 foci and unrepaired or slowly repaired recombination intermediates. These observations are consistent with the role suggested for Rad54 in yeast, namely its ability to remove Rad51 filaments from dsDNA (41). Possibly, RAD54B has also a role in meiotic HR, because when both RAD54 and RAD54B are absent compared to only RAD54, a slight increase in abnormal RAD51 distribution is detected. Altogether, while RAD54 and RAD54B contribute to HR, their mechanisms in mammalian meiosis are still unclear.

Here, we determined the cryo-electron microscopy (cryo-EM) structures of the complexes between P-motifs of BRCA2 and RAD54B and ssDNA-DMC1 filaments at high resolution. We uncovered how these P-motifs specifically recognize the meiotic recombinogenic filaments, resulting in an increase in their stability.

## Material and methods

### Protein Expression and Purification

DMC1 and its variants were produced and purified as described before (30). Briefly, constructs for bacterial expression of human his-TEVsite-DMC1 (wild-type and variants) were transformed into Rosetta2 DE3 pLysS *Escherichia coli* bacteria. The bacteria were plated on selective (50 µg/mL kanamycin, 30 µg/mL chloramphenicol) LB-agar plates. Colonies were used to inoculate 200 ml selective LB media; the culture was grown overnight at 37 °C with shaking and protein expression was induced using 0.5 mM IPTG. Cells were cultured for another 3 hours at 37 °C, collected by centrifugation and frozen. The pellet was thawed in the following buffer: 3M NaCl, 100 mM Tris pH 7.5, 10% glycerol, 0.5 mM EDTA, 5 mM ß-mercaptoethanol, protease inhibitors (Roche), and sonicated on ice. DMC1 was purified by affinity chromatography using a 5 ml HisTrap Fast Flow column (Cytiva), the eluted fraction was supplemented with the TEV protease and the sample was dialyzed overnight at 4 °C. It was loaded on a 5 ml HiTrap heparine column (Cytiva) and then a 5 ml Capto Q column (Cytiva). The protein was detected at about 500 mM NaCl in 50 mM Tris pH 8.0, 10% glycerol and 1 mM DTT, and further dialysed in 50 mM Tris pH 8.0, 150 mM NaCl and 1 mM DTT before characterization by biochemical and biophysical methods. RAD51 was expressed and purified using a similar protocol, and also finally dialysed in 50 mM Tris pH 8.0, 150 mM NaCl and 1 mM DTT.

Untagged wild-type human RAD51 was purified as described before (42). The RAD51 F86E variant was expressed from pET11M-His6-TEV-hRAD51 as N-terminal His6-tagged protein in E. coli Rosetta 2(DE3) cells (Merck) cultured in 6.4 L LB medium at 30°C overnight. The cells were collected, re-suspended in 120 mL buffer D (50 mM Tris-HCl (pH 8.0), 10% glycerol, 500 mM NaCl, 10 mM imidazole, 2 mM dithiothreitol), and lysed by sonication. The supernatant was then separated from the cell debris by centrifugation (39, 191g) for 30 minutes at 4°C and mixed with HisPur Ni-NTA resin (2 mL; Thermo Fisher Scientific) at 4°C for 1 hour. The beads were packed into a disposable chromatography column and were washed with 400 mL buffer D. His6-tagged RAD51 was eluted with 100 mL buffer D containing 500 mM imidazole in 4-mL fractions. Fractions containing the His6-tagged RAD51 protein were collected, and the concentration was measured by the Bradford method (ref, Bradford paper). The His6 tag was cleaved with TEV protease (50 μg per 1 mg eluate) during dialysis against 4 L buffer E (50 mM Tris-HCl (pH 8.0), 10% glycerol, 0.25 mM EDTA, 50 mM KCl, 2 mM dithiothreitol) overnight. The protein was then loaded onto a HiTrap Heparin HP column (1 mL; Cytiva) pre-equilibrated with buffer E, washed with 10 mL buffer E, and eluted with 10 mL buffer E containing 600 mM KCl. Fractions containing RAD51 were pooled and applied to Superdex200 16/60 pg pre-equilibrated with buffer E. Monomeric RAD51 was collected and subsequently applied to Capto Hires Q (1 mL) equilibrated with buffer E. The column was washed with 10 mL buffer E and eluted with 10 mL buffer E containing 600 mM KCl. Eluted RAD51 was collected and dialyzed against 1 L buffer G (20 mM HEPES-NaOH (pH 7.4), 10% glycerol, 150 mM NaCl, 0.1 mM EDTA, 2 mM dithiothreitol), snapfrozen, and stored at −80 °C.

### Cryo-Electron Microscopy

For cryo-EM experiments, a biotinylated capped ssDNA substrate was first prepared by incubating 5 μM monomeric streptavidin with 1 μM (in molecule) of an 80-nt ssDNA biotinylated at both ends, in a buffer containing 20 mM Tris-HCl pH 8 during 30 min at room temperature. Then, DMC1 filaments were assembled by incubating 40 μM (in nucleotides) of this biotin-streptavidin capped ssDNA substrate with 13.4 µM DMC1 in a buffer containing 10 mM Tris-HCL pH 8, 50 mM NaCl, 3 mM CaCl_2_, 1 mM DTT and 2 mM AMP-PNP during 20 min at 37 °C. To prepare cryo-EM grids, 3 μL of the reaction were applied to a plasma-cleaned (15 mA, 30 s) 200-mesh gold carbon grid. Grids were then blotted and plunged into liquid ethane using a Vitrobot Mark IV (FEI) with a blotting force of 0 and a blotting time of 3 s. For the DMC1 filaments bound to BRCAP-PL, 1 µl of peptide was added to the cryo-EM grids already loaded with the filaments (final ratio: 2 peptides / 1 DMC1), and then, after 3 min, the grids were blotted. For the DMC1 filaments bound to RAD54B-PL, 1 µl of peptide was added to the sample preparation during 5 min at 37 °C before deposition on the grids (final ratio: 2 peptides / 1 DMC1).

For the analysis of the ssDNA-DMC1 grids, a total of 8, 968 movies were processed in real time using CryoSPARC Live (Structura Biotechnology) and subsequent image processing was performed in CryoSPARC v4.7.1 (43). Movies were corrected with Patch Motion Correction and Patch CTF Estimation. Approximately 1, 900, 00 particles were initially picked using a combination of template-free Blob Picker and Filament Tracer. The particles were extracted with a box size of 336 pixels for initial processing. Following 2D classification, about 580, 000 particles were subjected to Ab Initio 3D Reconstruction. The resulting maps were used as input for Heterogeneous Refinement and 3D Classification jobs. In total, 267, 809 particles and the best 3D map were used for Reference Motion Correction and Non-Uniform Refinement, yielding a 2.16 Å reconstruction.

For BRCA2-PL bound to ssDNA-DMC1 filaments, a total of 6884 movies were processed in CryoSPARC Live. They were corrected with Patch Motion Correction and Patch CTF Estimation. Approximately 2, 000, 000 particles were initially picked using a combination of template-free Blob Picker and Filament Tracer. The particles were extracted with a box size of 336 pixels for initial processing. Following 2D classification, about 590, 000 particles were subjected to Ab Initio 3D Reconstruction. The resulting maps were used as input for Heterogeneous Refinement and 3D Classification jobs. In total, 197, 574 particles and the best 3D map were used for Reference Motion Correction and Non-Uniform Refinement, yielding a 1.90 Å reconstruction.

For RAD54B-PL bound to ssDNA-DMC1 filaments, a total of 10, 228 movies were imported into cryoSPARC. They were corrected with Patch Motion Correction and Patch CTF Estimation. Approximately 1, 500, 000 particles were initially picked using a combination of Blob Picker and Template Picker. These particles were extracted with a box size of 336 pixels. Following 2D classification, about 1, 300, 000 particles were re-extracted with the same box size. These particles were subjected to Ab Initio 3D Reconstruction. The resulting maps were used as input for Heterogeneous Refinement and 3D Classification jobs. In total, 409, 270 particles and the best 3D map were used for Reference Motion Correction and Non-Uniform Refinement, yielding a 2.04 Å reconstruction.

For all datasets, the processing was performed in cryoSPARC version 4.5.0.1 (43) and the maps were visualized using UCSF ChimeraX 1.10.1 (44).

For 3D reconstructions, initial models were calculated using ModelAngelo version 1.0.13 (45). These models were manually corrected using Coot version 0.9.8.96 (46) and refined with Servalcat version 0.4.126 (47) using the helical reconstruction mode (servalcat refine_spa_norefmac --pg C1 --twist 55.7 --rise 15.92) defining the asymmetric unit as one monomer of DMC1, plus 3 nucleotides, the AMP-PNP and the peptide if present.

### GST Pull-Down Assays

GST pull-down assays were performed as described previously (30). Bacterial expression constructs for expression of his-GST-TEV-tagged RAD54B fragments and their amino acid substitution variants were engineered using Gibson assembly in pETM-30 vector and sequence-verified. Constructs were transformed into Rosetta2 DE3 pLysS *E. coli* expression strain. Two mL of selective LB (50 µg/mL kanamycin, 30 µg/mL chloramphenicol) media was inoculated, the culture was grown overnight at 37 °C with shaking, diluted to 20 mL, further incubated till OD_600_ reached 0.6-0.8, induced with 0.2 mM IPTG, grown for additional 3 h at 37°.

Cells were collected by centrifugation, pellet was resuspended in 1 mL NETT+D buffer (NaCl 100 mM, 50 mM Tris-HCl pH 7.5, 5 mM EDTA pH 8, Triton X100 0.5%, freshly supplemented with protease inhibitors (Roche), 1 mM DTT) and sonicated (10× 5 sec on, 10 sec off). Lysate was transferred to Eppendorf minicentrifuge tubes and cleared by centrifugation in (30 min at 4 °C). Supernatant was mixed with 20 µL GSH-Sepharose beads (GE Healthcare 17-5132-01) and incubated overnight at 4 °C. Beads were collected by centrifugation (500 rcf, 2 min, 4 °C), washed with NETT+ buffer ((NaCl 100 mM, Tris-HCl pH 7.5 50 mM, EDTA pH 8 5 mM, Triton X100 0.5%, freshly supplemented with protease inhibitors (Roche)), and incubated with DMC1 protein solution (∼2 µg in 1 mL NETT+) for 1.5 h at 4 °C. Prior to incubation, bead suspension was vortexed, a 40 µL aliquot was collected as input, mixed with 12 µL 5× sample buffer (50% glycerol, 250 mM Tris HCl pH 6.8, 10% SDS, 0.5% bromophenol blue, 0.5 M β-mercaptoethanol) and denatured for 5 min at 95 °C. After incubation with DMC1, beads were washed three times with NETT+ buffer and incubated with 25 µL of Laemmli sample buffer for 5 min at 95 °C to elute bound proteins. Samples (5 µL of eluate) were run on a 13% SDS-PAGE, transferred to PVDF membrane, blocked (5% skim milk in PBS-T (PBS with 0.05% Tween-20)) and immunodetected with a mixture of rabbit anti-RAD51 pAb (1:20000 (61)) and mouse anti-GST mAb (1:5000, B-14 Santa Cruz, sc-138) antibodies overnight. After washes, the membranes were incubated with fluorescently labelled secondary antibodies (anti-mouse CF680 (Sigma #SAB460199), anti-rabbit CF770 (Sigma #SAB460215), washed (5 × 5 min PBS-T) and scanned using Odyssey CLx imaging system (LI-COR).

### Nano-differential scanning fluorimetry (nanoDSF)

The thermal stability of DMC1 protein variants (WT, F89A, D175K, D180K, E213K and Δ18) was assessed using a Prometheus Panta instrument (NanoTemper Technologies, Munich, Germany) employing nano-differential scanning fluorimetry (nanoDSF). Protein samples were diluted to a final concentration of 0.5-0.75 mg/mL in a buffer containing 25 mM Tris-HCl (pH 7.5), 100 mM NaCl, and 2 mM DTT. Samples were loaded into standard nanoDSF capillaries (NanoTemper Technologies) according to the manufacturer’s instructions. Thermal unfolding experiments were performed by increasing the temperature from 25 °C to 95 °C at a heating rate of 1 °C/min. Intrinsic protein fluorescence was monitored simultaneously at 330 nm and 350 nm, and the fluorescence ratio (F350/F330) was calculated throughout the temperature ramp. The thermal unfolding profiles shown correspond to the first derivative of the fluorescence ratio. Protein melting temperatures (Tm) were determined from the minima of the first derivative of the F350/F330 ratio as a function of temperature. Data acquisition and analysis were performed using the PR.Control and PR.ThermControl software packages (NanoTemper Technologies).

### BioLayer Interferometry (BLI)

Interactions between DMC1 (2 µM) and biotinylated peptides — BRCA2-P (Biotin-GSG-TTGRPTKVFVPPFKTKSHFH-NH₂), RAD54B-P (Biotin-GSG-GNSFKKPKFIPPGRSNPG-NH₂), RAD54B-P F20A (Biotin-GSG-GNSFKKPKAIPPGRSNPG-NH₂), FIGNL1 P-motif (Biotin-GSG-RSRGILGKFVPPIPKQDGGE-NH₂), FIGNL1 A-motif (Biotin-GSG-SSLPTFKTAKEQLWVDQQKK-NH₂), and RAD51AP1 P-motif (Biotin-GSG-SAESKKPKWVPPAASGGSRS-NH₂), synthesized by Proteogenix and Genecust, were measured by biolayer interferometry (BLI) using an Octet RED96 instrument (FortéBio). Experiments were performed at 25°C. Streptavidin (SA) biosensors were hydrated for 20 min in buffer containing 25 mM Tris-HCl (pH 7.5), 100 mM NaCl, and 5 mM β-mercaptoethanol. No non-specific binding of DMC1 to unloaded SA biosensors was detected. Biotinylated peptides (0.25 µg/mL) were immobilized onto SA biosensors and washed in buffer prior to the association step. Association was monitored by incubating the sensors with DMC1 for 300 s, and dissociation was measured by transferring the sensors into buffer (25 mM Tris-HCl, 100 mM NaCl, 5 mM β-mercaptoethanol, 0.05% Tween-20) for 600 s.

The interaction between DMC1 (WT or mutants) filaments assembled on a biotinylated 100-nt ssDNA (Biotin-AATTCTCATTTTACTTACCGGACGCTATTAGCAGTGGGTGAGCAAAAACAGGAAGGCAAAATGCCGCAAAAAAGGGAATAAGGGCGACACGGAAATGTTG; synthesized by IDT, Integrated DNA Technologies) and peptides (BRCA2-P, RAD54B-P, and RAD54B-P K17E or K19E) was also analyzed by BLI. Streptavidin (SA) biosensors were hydrated in buffer containing 10 mM Tris-HCl (pH 8.0), 100 mM NaCl, and 5 mM β-mercaptoethanol. Biotinylated ssDNA (60 nM) was immobilized onto SA biosensors, which were then incubated with DMC1 (WT or mutants, 1 µM) in filament assembly buffer (10 mM Tris-HCl, pH 8.0, 100 mM NaCl, 5 mM β-mercaptoethanol, 2 mM CaCl₂, 2 mM ATP, and 0.05% Tween-20) until a signal of ∼2.5 nm was reached. Following a dissociation step in the same buffer to assess filament stability and re-establish baseline, binding kinetics were measured using increasing concentrations of peptides prepared by twofold serial dilution (30, 15, 7.5, 3.75, and 1.87 µM). Apparent Kd values were determined using a steady-state analysis.

### Isothermal titration calorimetry (ITC)

The interaction between RAD54B-P and the DMC1 octamer was characterized by isothermal titration calorimetry (ITC) using a VP-ITC calorimeter (GE Healthcare). Purified DMC1 WT protein and synthetic peptide (Genecust)—RAD54B-P (Ac-^12^GNSFKKPKFIPPGRSNPG^29^-NH₂) were used. The buffer consisted of 25 mM Tris-HCl (pH 7.5), 100 mM NaCl, and 5 mM β-mercaptoethanol. Experiments were performed at 20 °C. Proteins in the cell (8–17 µM) were titrated with peptide solutions (80–170 µM) loaded into the syringe, using 12 µL injections at 240 s intervals. Data were acquired and analyzed with Origin 7.0 software supplied by the manufacturer.

### Transmission Electron Microscopy

Negative stain transmission electron microscopy was used for measuring the ssDNA-DMC1 filament lengths: DMC1 filaments were first assembled by incubating 3 mM in nucleotides of a 100-nt ssDNA with 1 μM DMC1 (1 protein per 3 nt) in a buffer containing 10 mM Tris-HCl pH8, 50 mM NaCl, 3 mM CaCl_2_, 2 mM ATP and 1 mM DTT, 20 minutes at 37 °C. Then, 0 or 625 nM of either BRCA2-PL or RAD54B-PL peptide were added to the reaction during 5 minutes at 37 °C. A drop of the reaction was directly deposited on a carboned copper grid pre-activated with a glow discharge (plasma) and then observed in bright-field using a Zeiss 902 transmission electron microscope. Images were captured at a magnification of 85, 000× with a Veleta CCD camera and analyzed with iTEM software (both Olympus Soft Imaging Solution). For the quantifications, the length of filaments was measured on at least 2 independent experiments with a total of at least 200 molecules measured.

### Electrophoretic Mobility Shift and D-loop Assays

In EMSA, DMC1 filaments were formed as in the TEM experiments, by incubating 3 mM a 100-nt ssDNA (CCTACATACCTCGCTCTGCTAATCCTGTTACCAGTGGCTGCTGCCAGTGGCGATAAGTCGTGTCTTACCGGGTTGGACTCAAGACGATAGTTACCGGAT) labeled with Cy5 with 1 μM DMC1 in a buffer containing 10 mM Tris-HCl pH7.5, 50 mM NaCl, 3 mM CaCl_2_, 2 mM ATP and 1 mM DTT, 20 minutes at 37°C. 0, 0.25, 0.5, 0.75 or 1 μM BRCA2-PL or RAD54B-PL peptide were then added to the reaction 5 minutes at 37 °C. Protein–DNA complexes were fixed with 0.01 % glutaraldehyde 5 minutes at room temperature. The reaction products were then analyzed using 0.75 % agarose gel in 0, 5x Tris acetate EDTA, run 30 minutes at 80V, 4 ◦C. Images were acquired using a Typhoon imager (GE Healthcare Life Science).

For D-loop assays, DMC1 filaments were formed in same conditions as for EMSA analysis, using same incubation conditions and buffer. In the second step, 15 nM in molecules of homologous dsDNA donor (pUC19 plasmid purified on MiniQ ion exchange chromatography column) were introduced in the reaction and the mix was incubated during 30 minutes at 37°C. Then the reaction was stopped and deproteinized using 0.5 mg/mL Proteinase K, 1% SDS, 12.5 mM EDTA and incubated 30 minutes at 37°C. A 1% TAE agarose gel was run at 80 V, for 30 minutes. The reaction products were analyzed using a Typhoon imager (GE Healthcare Life Science). D-loop yield was quantified on Image J software as a percentage of the D-loop product relative to the total DNA.

For the EMSA assays on DMC1 ΔN18 and ΔN81, 3 µM nucleotides (nt) (30 nM molecules) of single-stranded DNA (Cy5-TGCTTCCGGCTCGTATGTTGTGTGGAATTGTGAGCGGATAACAATTTCACACAGGAAACAGCTATGACCATGATTACGAATTCGAGCTCGGTACCCGGGG-3ʹ) were incubated with DMC1 WT or mutant proteins (1 µM) for 10 min at 37 °C in buffer containing 10 mM Tris-HCl (pH 7.5), 50 mM NaCl, 2 mM CaCl₂, 2 mM ATP, and 1 mM DTT. Protein–DNA complexes were crosslinked by an addition of 0.01% glutaraldehyde for 5 min at room temperature. Reaction products were resolved on a 0.75% agarose gel in 0.5× Tris-acetate-EDTA (TAE) buffer at 80 V and 4 °C for 30 min. Gels were imaged using a Chemidoc system (Bio-Rad) with Cy5 detection.

### AlphaFold Calculations

The AlphaFold 3 webserver was used to calculate models for ssDNA-RAD51(2x)-BRCA2-PL and ssDNA-RAD51(2x)-RAD54B-PL complexes. Five models were obtained for each complex. The models were visualized using UCSF ChimeraX 1.10.1 (44). A representative model of each complex is shown in Figure S7.

### Molecular Modelling and MD simulations

To achieve Molecular Dynamics (MD) simulation in explicit solvent, the first step of the work consisted in building the starting molecular model, which was based on the cryo-EM structure of ssDNA-DMC1-BRCA2-PL. This structure permitted to define the cartesian coordinates of residues 20 to 329 for 6 DMC1 chains (A, B, C, D, E, and F). It also contains the coordinates of the atoms of the single strand DNA and of a short region of the BRCA2 peptide (residues 2403 to 2412). The starting coordinates of the residues not visible in the cryo-EM structure were further rebuilt, i.e. residues 1-19 located in N-terminal position of the DMC1 protomers and the BRCA2-PL residues except for residues 2403 to 2412. We generated initial coordinates of the atoms of the DMC1 protomers and of the BRCA2-PL peptide not visible in the cryo-EM structure using Modeller 10.6 (48). We kept only one BRCA2-PL peptide in interaction with chain C of the DMC1 filament (segid DMCC). This choice allows to define possible interactions of the distal parts of the peptide without interference of other BRCA2-PL peptides during the simulation. Structural models were analysed using in house CHARMM scripts and UCSF ChimeraX 1.12 (44).

The initial ssDNA-DMC1-BRCA2-PL structure was refined using successive steps of energy minimization and restrained molecular dynamics with CHARMM v49b1 (49), using the CHARMM36m force field (50). The system was solvated in a TIP3P water cubic box using the solvate module of VMD 1.9.4 (51) with a minimum distance of 12 Å between the protein/peptide and the box boundaries in all directions. The system was subsequently neutralized and supplemented with Na⁺ and Cl⁻ ions to reach an ionic strength of 150 mM. Energy minimization was performed for 10, 000 steps under positional restraints to maintain the structure close to the initial conformation. Several minimization cycles were applied, during which the force constant on atomic positions was gradually reduced from 10 kcal·mol⁻¹·Å⁻² to 0. This step was followed by a 2 ns molecular dynamics equilibration step in the NVT ensemble. Production simulations were then carried out in the NPT ensemble. To account for stochastic effects, three MD trajectories were run with different initial velocities for a total duration of 1 µs. Temperature was maintained at 310 K and pressure at 1 atm using Langevin dynamics. During production, a time step of 2 fs was used, and multiple time stepping was applied using the rRESPA algorithm, with long-range electrostatic interactions computed every 4 fs using the Particle Mesh Ewald (PME) method with a real-space grid spacing of 1 Å. Periodic boundary conditions were applied. A cutoff distance of 12 Å was used for short-range electrostatic and van der Waals interactions. Molecular dynamics simulations were performed using NAMD 3.0.1 (CUDA version) (52). Trajectory analyses were conducted using in-house CHARMM scripts. MD trajectories were analysed using ChimeraX 1.12 and VMD. Figures were generated with ChimeraX 1.12 and Gnuplot 6.0 (53).

## Results

### A BRCA2 fragment of 10 residues containing the PhePP motif stably interacts with the ssDNA-DMC1 filaments

We and others previously found that a BRCA2 peptide containing the PhePP motif (28) binds and stabilizes ssDNA-DMC1 filaments, but at the structural level, the interaction between BRCA2 (either T2398-H2417 or R2401-S2414) and DMC1 was only studied by X-ray crystallography in the absence of DNA (30, 31). To address this, we structurally characterized the complex between a larger BRCA2 fragment BRCA2-PL (G2379-Q2421) containing the PhePP motif and the ssDNA-DMC1 nucleoprotein filament using cryo-EM. First, we determined a high-resolution structure of the ssDNA-DMC1 filament without any peptide at a global resolution of 2.2 Å. Then, we similarly characterized the BRCA2-PL-bound filament. We observed that the BRCA2 peptide caused filament bundling and aggregation. Therefore, ssDNA-DMC1 filaments were first deposited on the cryo-EM grids; the BRCA2 peptide was then added before blotting and freezing. Processing of the cryo-EM images produced a 3D reconstruction of the ssDNA-DMC1-BRCA2-PL complex at a global resolution of 1.9 Å (**Figures S1A-B; Table S1**). Both reconstructions revealed densities corresponding to the ssDNA-DMC1 filament (**Figure 1A, left panel**). The densities corresponding to the ATPase domain of the recombinases are the best defined, whereas those of the solvent exposed regions of the N-terminal folded domain of DMC1 are the least resolved; no density is observed for the 19 N-terminal residues of DMC1 and 5 residues of the ATPase loop 2 (**Figures S1A-D**). The 3D structure of the ssDNA-DMC1 filament bound to BRCA2-PL is superimposable with that of the filament without peptide, as solved in our conditions (RMSD on the Cα atoms of a DMC1 protomer: 0.57 Å) and as previously published (PDB 7C9C; RMSD on the Cα atoms of a DMC1 protomer: 0.77 Å). In our first reconstruction, the BRCA2 peptide is clearly visible, its central region being defined at a resolution between 2.0 and 2.5 Å (**Figure 1A, right panels; Figure S1A**). It contacts the DMC1 ATPase domain, in a region that is located at the interface with both the N-terminal region of the same DMC1 protomer and the ATPase domain of the adjacent DMC1 protomer. Within the BRCA2 peptide, 10 residues, from T2403 to T2412, could be robustly positioned in the cryo-EM density. This region corresponds to the well-conserved motif of the BRCA2 peptide (**Figure 1B**). A large region at the N-terminus of the peptide (G2379-P2402) could not be detected (**Figure 1A, right lower panel**), consistent with low evolutionary conservation. Careful analysis of the density corresponding to the peptide revealed that, while residues T2403 to P2409 (including the F-V-P-P motif) adopt a single conformation, residues F2410-T2412 exhibit two alternative conformations (**Figure 1C**), indicative of residual dynamics in their bound state.

**Figure 1.**
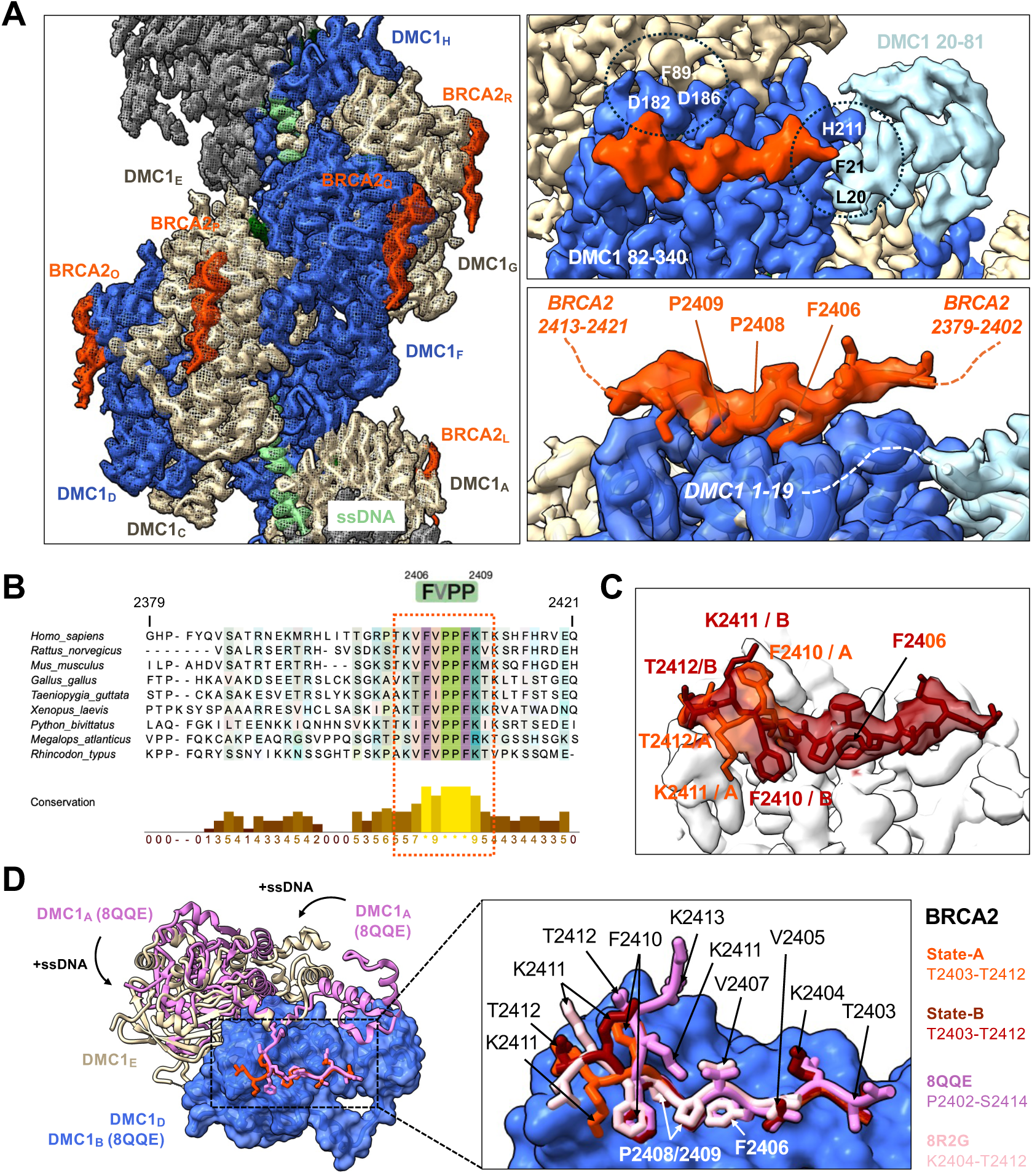
The PhePP BRCA2 region G2379-Q2421 (BRCA2-PL) interacts with the ssDNA-DMC1 filament through a stable core sequence of 10 conserved residues centered on the FVPP motif. **(A)** Cryo-EM map of the ssDNA-DMC1 filament (AMP-PNP and Ca^2+^) in the presence of the 43-aa BRCA2-PL peptide (resolution 1.94 Å; Figure S1; Table S1). The ssDNA, 8 DMC1 monomers and 8 BRCA2 peptides were docked into the map, and the map was colored as the fitted structure. Two zoom views are shown in the right panels. The upper zoom view highlights that the BRCA2 peptide is bound in close proximity with the interfaces between (i) the folded N-terminal (light blue) and ATPase (royal blue) domains of a DMC1 protomer, as well as (ii) the ATPase domain of this protomer (royal blue) and the linker region (82–98) of the adjacent protomer (wheat). Residues involved in these interfaces are marked. The lower zoom view stresses that, within the 43-aa BRCA2 fragment, only 10 residues stably interact with the ATPase domain of DMC1 (buried surface: 880 Å^2^), and, within the DMC1, the region stably assembled onto ssDNA begins at residue L20. **(B)** Sequence alignment of BRCA2-PL within vertebrates produced using Jalview 2.11.5.1. The core sequence stably bound to ssDNA-DMC1 as observed by cryo-EM is marked by a dotted red box. **(C)** Two alternative BRCA2-PL structures fitted into the cryo-EM map. Even in the stably bound core of the peptide, two different conformations were detected for residues F2410, K2411 and T2412 (orange red versus dark red). **(D)** Comparison of the BRCA2 conformations bound to the ssDNA-DMC1 filament versus the DMC1 octamer (8QQE: full-length DMC1; 8R2G: DMC1 deleted from its N-terminal domain), showing that the fragment T2403-P2409 has a similar structure in all contexts, whereas the conformation of F2410-T2412 varies. In the left panel, the cryo-EM structure of the filament with BRCA2-PL in state A is compared to the crystal structure of oligomeric full-length DMC1 with the 20-aa BRCA2-P peptide. A zoom view of the peptide conformations is shown on the right panel. The recombinase structures are displayed as cartoons and the DMC1 chain D surface is shown in royal blue. The BRCA2 peptides are shown as cartoons, with their side chains in sticks.

We compared the conformation of the BRCA2 peptide bound to either octameric DMC1 (30, 31) or DMC1 assembled into a nucleoprotein filament, as reported in this study. We previously observed that, upon binding of the BRCA2 peptide T2398-H2417 (BRCA2-P) to the DMC1 octamer, a 13-aa BRCA2 sequence was stably positioned onto a single DMC1 protomer and the N-terminal folded domain of the adjacent DMC1 was partially docked against the ATPase domain of the BRCA2-bound protomer (30). The structure of the DMC1 oligomer is modified when the recombinase binds to ATP and is loaded onto ssDNA: ATP is captured at the interface between two protomers and the relative positioning of the protomers shifts (19). The position of its N-terminal domain changes too, with it becoming docked between two different protomers in the filament. Nevertheless, the BRCA2 residues T2403-P2409 bind at the same DMC1 P-site and adopt the same bound conformations (**Figure 1D**). Only the positions of residues F2410-T2412 vary between the crystal and the cryo-EM structures. These residues already showed alternative conformations in the cryo-EM filament structure. Altogether, the BRCA2 peptide exhibits a core binding motif with a single bound state, as well as additional residues that have different bound states in solution as well as depending on the structural context.

### When comparing BRCA2 TR2 and PhePP, only the central motif F-x-x-P bind to a surface conserved between RAD51 and DMC1

In BRCA2, two different motifs bind to either ssDNA-RAD51 or ssDNA-DMC1: the TR2 motif binds to RAD51 filaments whereas the PhePP motif targets DMC1 filaments. We asked how BRCA2 regions specifically recognize different presynaptic filaments. Therefore, we compared our BRCA2 PhePP-DMC1 structure to the previously reported BRCA2 TR2-RAD51 structure (32) Both filament structures can be closely superimposed, as published (19, 20) (**Figure S2A**). When zooming on the bound peptides, the TR2 density is clearly visible for BRCA2 A3284-C3304 (EMDB 17584). The structures of PhePP T2403-T2412 and TR2 A3284-C3304 overlap in their central parts, with the side chains of F2406 and P2409 being buried in the same cavities as F3298 and P3301, respectively (**Figure 2A**). Moreover, the residues forming these cavities are conserved between DMC1 and RAD51. We also compared the BRCA2 PhePP-DMC1 binding interface to the RAD51AP1 binding interface with RAD51 presynaptic filaments (6). In this last structure, the C-terminus of RAD51AP1 (Q330-A349) overlaps only locally with PhePP, but the side chains of L345 and H346 are still buried in the same cavities as F2406 and P2409, respectively (**Figure S2B**). Thus, while BRCA2 PhePP shares a F-x-P-P motif with only BRCA2 TR2, all three BRCA2 and RAD51AP1 peptides exhibit a motif that docks at the center of the recombinase P-site, in a region that is conserved between DMC1 and RAD51 (**Figure 2A; Figure S2A**). Outside of these motifs, the peptides interact with residues that are not conserved between the recombinases (**Figure 2A; Figure S2A**). We propose that such contacts are responsible for the specific recognition of DMC1 versus RAD51 filaments.

**Figure 2.**
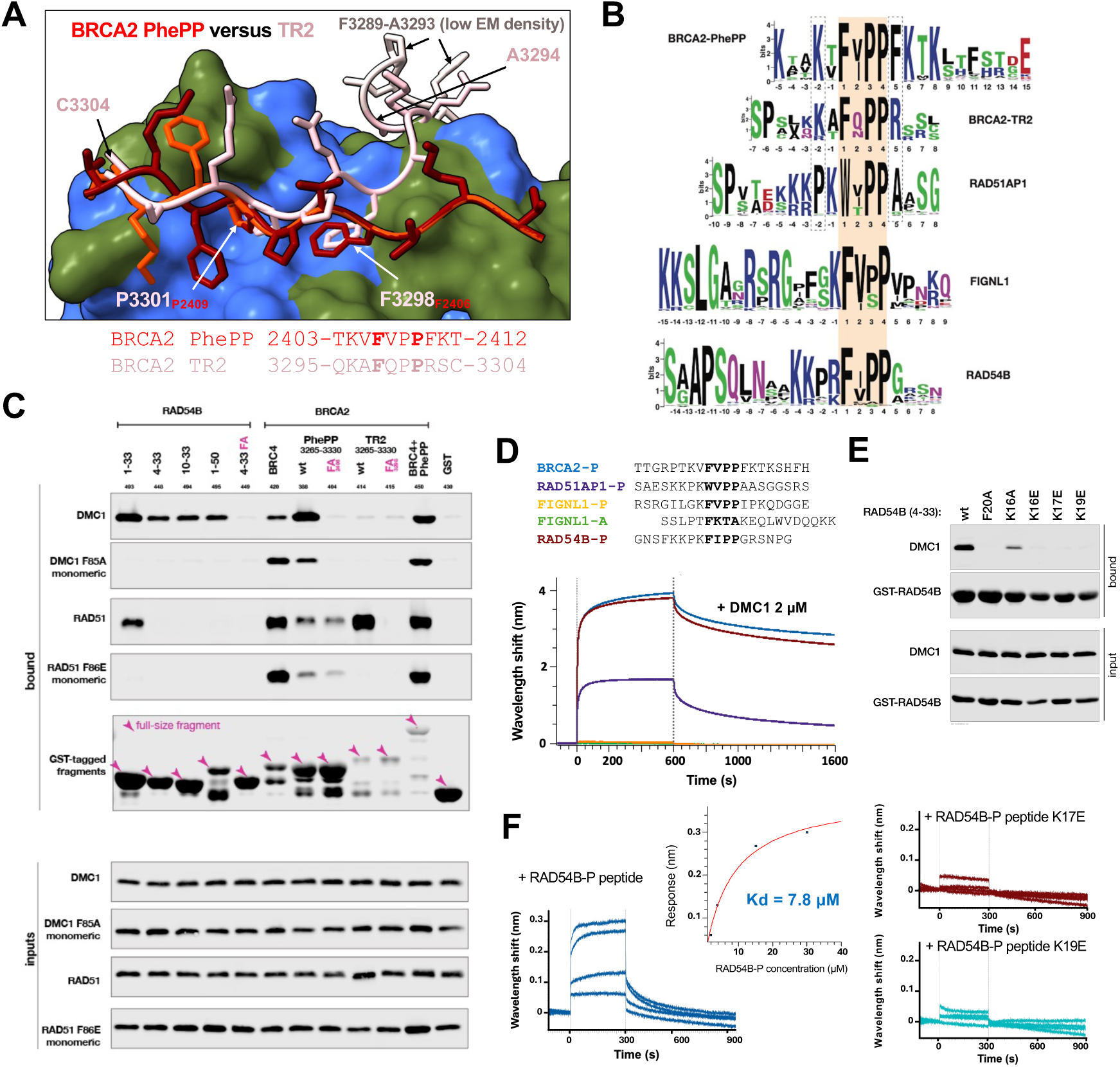
The P-motif shared by BRCA2 PhePP and TR2 is also found at the N-terminus of the DNA translocase RAD54B. **(A)** Comparison of the BRCA2 PhePP structures bound to the ssDNA-DMC1 filament and the BRCA2 TR2 structure bound to the ssDNA-RAD51 filament (8PBC; ATP and Ca^2+^). The DMC1 surface is shown in royal blue, except for residues not conserved in RAD51 that are in dark green. The RAD51 protomer is omitted for clarity. The BRCA2 peptides are shown as cartoons, with their side chains in sticks. **(B)** Search for proteins sharing P-motifs in HR proteins (WebLogo format). **(C)** GST-pulldown experiments performed using tagged peptides containing P-motifs (WT and variants). DMC1 (WT and F85A) and RAD51 were detected by immunoblotting. The RAD54B peptide variant FA corresponds to F20A. The PhePP and TR2 peptide mutants FA correspond to F2406A and F3298A, respectively, whereas the fused BRC4-PhePP peptide is indicated by BRC4+PhePP. **(D)** BLI experiment performed after attaching biotinylated peptides on SA biosensors and incubating these biosensors into solutions of DMC1. The sequences of the peptides are indicated on the top. Binding is observed between BRCA2-P, RAD51AP1-P, RAD54B-P and DMC1 octamers, whereas binding between the FIGNL1 peptides and DMC1 is too weak to be detected by BLI in these conditions. **(E)** GST-pulldown assays revealing binding of several RAD54B (4–33) wild-type and mutated peptides to purified DMC1. Not only F20, but also K16, K17 and K19, contribute to RAD54B binding to DMC1. **(F)** BLI experiment showing that RAD54B-P binds to ssDNA-DMC1 with an apparent Kd of 7.8 µM, whereas peptide mutants K17E and K19E show no detectable interaction with the DMC1 filaments.

### A P-motif binding to DMC1 is present at the N-terminus of RAD54B

We searched for additional disordered protein regions containing a conserved P-motif that binds to ssDNA-DMC1 filaments. We detected the FRBD region of FIGNL1 (fidgetin-like 1), known to interact with both RAD51 and DMC1 (30, 54). We also identified a new putative P-motif in the N-terminal region of the DNA translocase RAD54B (residues S4-N27) **(Figure 2B**). The binding properties of this RAD54B P-motif were characterized by GST-pulldown and compared to those of BRCA2 BRC4 (as an example of a BRCA2 A-motif), PhePP and TR2. Several RAD54B sequences containing the P-motif were tested (**Figure S3A**). The RAD54B-PXL (M1-E33) and RAD54B-PM (S4-E33) both bind to DMC1 wild-type, but no interaction with monomeric DMC1 F85A can be detected in our conditions (**Figure 2C; Figure S3B**). Moreover, only RAD54B-PXL binds to RAD51 in these same conditions, showing that the three first RAD54B residues are specifically essential for binding to RAD51. RAD54B-PXL binds to RAD51 wild-type, but not to monomeric RAD51 F86E (4). In line with previous reports, the BRC4 repeat of BRCA2 weakly binds to DMC1, either wild-type or monomeric, but this interaction is reinforced in both cases upon fusion of BRC4 to PhePP, which is consistent with BRC4 and PhePP binding to two distinct sites on DMC1 (30, 31). Finally, BRCA2 PhePP binds to DMC1 WT, but this interaction is reduced with monomeric DMC1 F85A, and it only weakly binds to RAD51, whereas BRCA2 TR2 specifically binds to RAD51. Thus, within the tested peptides, BRCA2 PhePP and RAD54B N-terminus preferentially bind to oligomeric DMC1, whereas BRCA2 BRC4 and TR2 preferentially bind to monomeric and oligomeric RAD51, respectively.

We compared the affinities of the different peptides for oligomeric DMC1 using BioLayer Interferometry (BLI). The RAD54B P-motif peptide G12-G29 (RAD54B-P) has a BLI interaction signal similar to that of BRCA2-P (**Figure 2D**) and a similar affinity for DMC1 as shown by ITC (**Figure S3C**), whereas the RAD51AP1 and FIGNL1 P-motif peptides have weaker BLI interaction signals, consistently with their lower affinity for DMC1, as reported previously (30). We further tested the role of the conserved aromatic residue of the RAD54B motif F-x-P-P for binding to DMC1: mutating F20 into alanine disrupts the interaction between RAD54B-PM and DMC1, as observed by GST-pulldown (**Figure 2E**). Similarly, RAD54B-P mutant F20A does not bind to DMC1, as revealed by BLI (**Figure S3C**). Finally, we tested the role of the conserved positively charged patch located just before the RAD54B motif FIPP (**Figure 2B**). We found that mutating K16, K17, and K19 into glutamic acid significantly decreases the DMC1 bound fraction detected by GST-pulldown (**Figure 2E**). Consistently, when measuring the apparent Kd of RAD54B-P for ssDNA-DMC1 filaments by BLI, we observed that mutating K17 and K19 into glutamic acid strongly disrupts binding (**Figure 2F**). Thus, both the positively charged patch K-K-P-K and the P-motif F-I-P-P contribute to the affinity of RAD54B for DMC1 assembled into presynaptic filaments.

### The 3D structures of DMC1 filament-bound BRCA2 and RAD54B are superimposable

BRCA2-P and RAD54B-P share a P-motif surrounded by positively charged residues, but their sequence similarity is restricted to F-x-P-P and their positively charged residues are mostly located at different positions (except for BRCA2 K2411 and RAD54B R25). We asked if they interact with the same acidic binding site in DMC1 (30). Therefore, we analyzed how the N-terminal region of RAD54B (RAD54B-PL: M1-G29) interacts with the recombinogenic ssDNA-DMC1 filament by cryo-EM. Processing of the cryo-EM images produced a 3D reconstruction of the ssDNA-DMC1-RAD54B-PL complexes at a global resolution of 2.0 Å (**Figure S4A; Table S1**). This reconstruction revealed densities corresponding to the ssDNA-DMC1 filament as well as the RAD54B-PL peptide (**Figure 3A; Figure S4B**). Here again, the DMC1 ATPase domain and the ssDNA are best defined, whereas the DMC1 N-terminal folded domain is less defined. The first 19 residues of DMC1, predicted as disordered, are not observed in the map (**Figure S4C**). Focusing on the peptide, the corresponding electron density is well-detected at a resolution of about 2.5 Å. The RAD54B residues P18-S26 could be confidently docked into this density. However, a large region at the N-terminus of the peptide (M1-K17) is not observed, despite the presence of highly conserved residues (**Figure 3B**).

**Figure 3.**
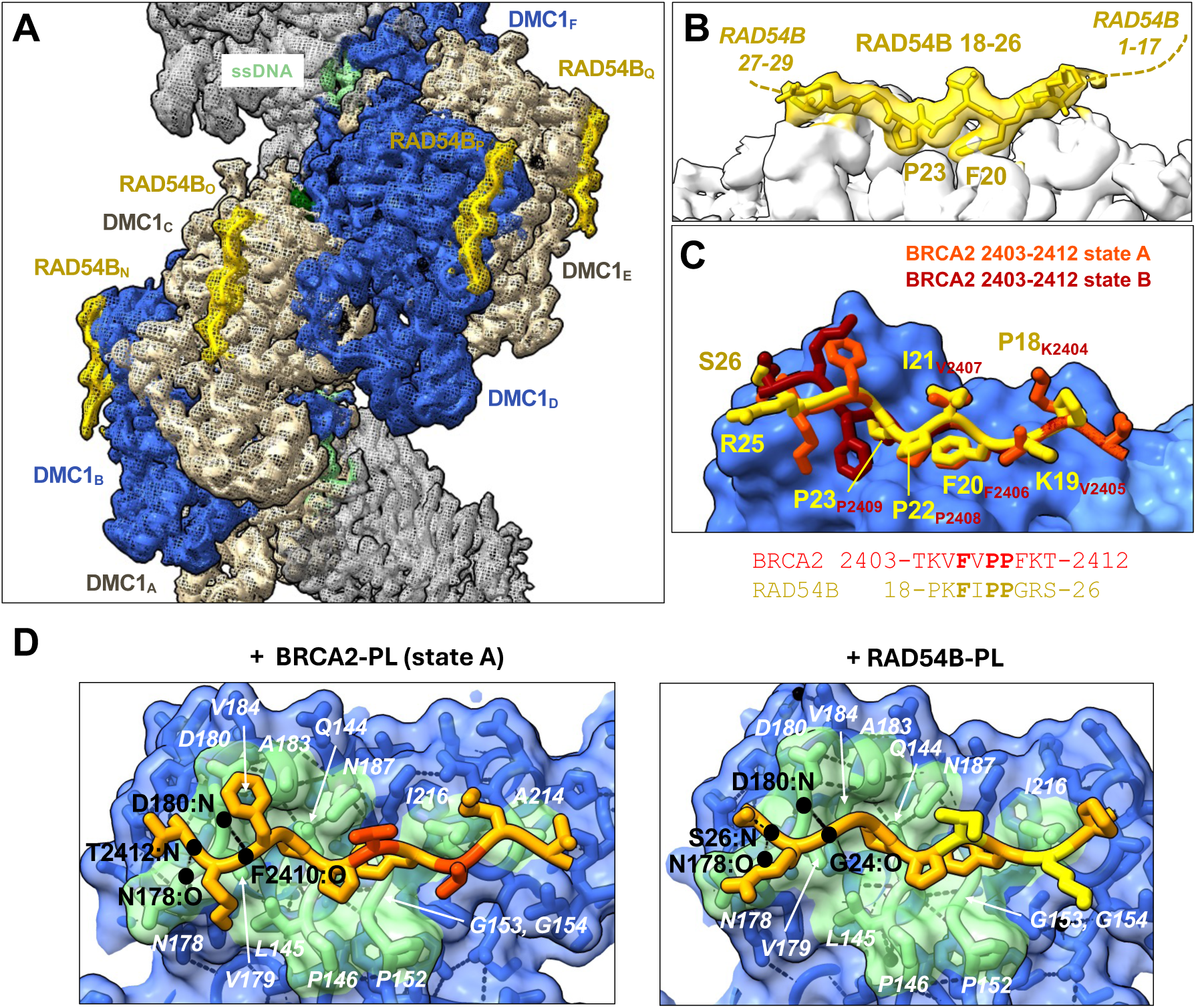
The BRCA2 and RAD54B peptides share the same bound structure and binding site on the ssDNA-DMC1 filaments. **(A)** Cryo-EM map of the ssDNA-DMC1 filament (AMP-PNP and Ca^2+^) in the presence of the N-terminal 29-aa RAD54B peptide (RAD54B-PL) (resolution 2.1 Å, see Figure S4 and Table S1). The ssDNA, 6 DMC1 monomers and 6 RAD54B-PL peptides were docked into the map, and the map was colored as the fitted structure. **(B)** Zoom view showing that 9 residues of the 29-aa RAD54B peptide stably interact with the ATPase domain of a DMC1 protomer (buried interface: 770 Å^2^). **(C)** Comparison of the BRCA2-PL and RAD54B-PL conformations bound to the ssDNA-DMC1 filament. The DMC1 chain D surface is shown in royal blue. The peptides are shown as cartoons, with their side chains in sticks. **(D)** Interfaces between DMC1 chain D (residues in contact with the peptides in light green) and the peptides (residues in contact with DMC1 in orange). Thirteen DMC1 residues form most of the peptide binding site: Q144, P146, P152, G153, G154, N178, V179, D180, A183, V184, N187, A214 (in front of BRCA2-PL T2403) and I216. Two backbone hydrogen bonds are formed between DMC1 and the peptides (in black dashed lines); the bound atoms are marked and labeled in black.

Superimposition of the cryo-EM structures of ssDNA-DMC1-BRCA2-PL and ssDNA-DMC1-RAD54B-PL revealed that fragments of similar sizes (9-10 residues) adopt a stable conformation when bound to ssDNA-DMC1. Moreover, these BRCA2 and RAD54B fragments adopt remarkably similar conformations (**Figure 3C**). The backbone conformation of RAD54B is particularly close to that of BRCA2 state A, and the side chains of the core sequences F-I/V-P-P exhibit superimposable structures. Both the BRCA2-PL and RAD54B-PL peptides contact twelve hydrophobic DMC1 residues that form a common binding groove, including two prolines, two glycines, an alanine, two valines, and an isoleucine (**Figure 3D**). They also interact with several polar DMC1 residues but form only two hydrogen bonds with DMC1; these hydrogen bonds are between backbone atoms and are conserved in both structures (**Figure 3D**). Unexpectedly, no stable salt bridge was identified between the positively charged residues of the peptides and the negatively charged residues of the DMC1 P-site. Altogether, our structural analysis showed that the interface between the P-motif containing peptides and the DMC1 filaments exhibits a large hydrophobic core and is stabilized by two hydrogen bonds that do not depend on the sequence of the ligands.

### The BRCA2 PhePP and RAD54B peptides stabilize DMC1 nucleoprotein filaments

We and others previously found that BRCA2 PhePP stabilizes the ssDNA-DMC1 filaments: addition of the peptide increases the length of the filaments (30) and protects the filaments from the disassembling effect of BRC4 (31). Here, we asked whether the N-terminal region of RAD54B could also stabilize the DMC1 nucleoprotein filaments. In conditions in which ssDNA-DMC1 filaments are assembled, as confirmed by negative-staining electron microscopy (EM), we found that the length of these filaments is multiplied by about 1.4 upon addition of 1 µM of the peptide BRCA2-PL or RAD54B-PL (**Figure 4A**). We also observed that D-loops are formed significantly more efficiently in the presence of either of these peptides (**Figure 4B**).

**Figure 4.**
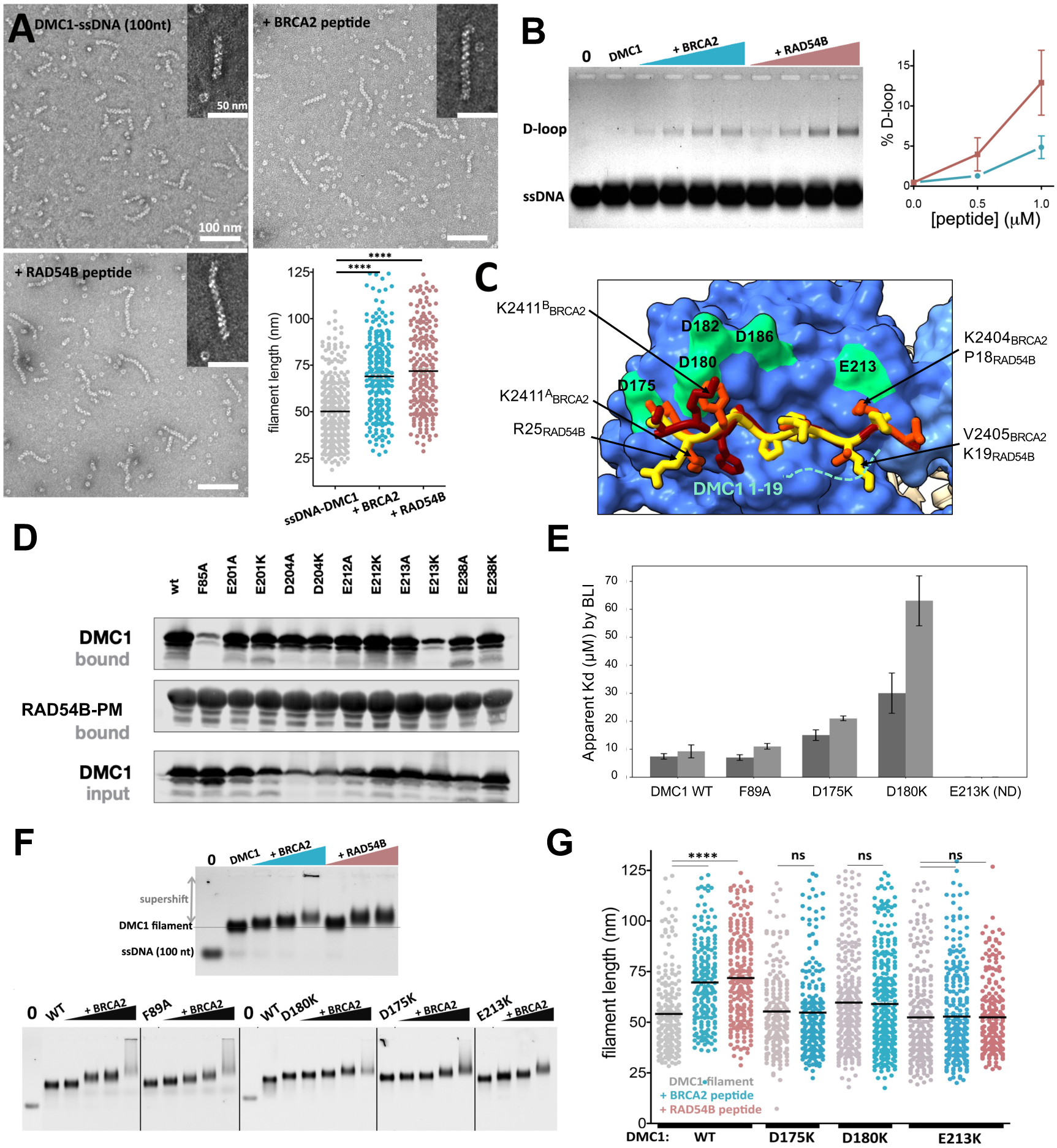
Both BRCA2-PL and RAD54B-PL stabilize ssDNA-DMC1 filaments and stimulate DMC1-mediated D-loop formation. **(A)** Quantification of ssDNA-DMC1 filaments lengths showing that BRCA2 and RAD54B peptides significantly promote and stabilize longer DMC1-filaments. Negative staining electron microscopy images of DMC1 filaments assembled on a 100-nt ssDNA in the absence and presence of BRCA2-PL or RAD54B-PL were obtained in conditions in which the peptides bind to the filaments, as confirmed by EMSA (see panel (F)). A total of 200 molecules were measured in two independent experiments. Filament lengths were plotted in the absence (grey) and presence of BRCA2-PL (blue) and RAD54B-PL (pink). **(B)** D-loop assays, in which filaments are assembled in absence or in presence of BRCA2 or RAD54B peptide, in the same conditions as in (B), and a pUC19 homologous donor plasmid is added to the reaction and incubated at 37° C during 30 minutes. The reaction is deproteinized before being run on an agarose gel. The D-loop yield was quantified in 2 to 3 independent reactions. **(C)** Superimposition of the cryo-EM structures of either BRCA2-PL or RAD54B-PL bound to ssDNA-DMC1. DMC1 is represented in surface mode, with its N-terminal region in cornflower blue and its ATPase domain in royal blue. The two BRCA2 alternative conformations are shown in orange red and maroon sticks, whereas the RAD54B peptide conformation is displayed in yellow sticks. DMC1 negatively charged residues in close proximity with the peptides are colored in green and labeled. Additionally, the negatively charged disordered N-terminus of DMC1 is displayed as a green dotted line. **(D)** GST-pulldown analysis of the interaction between GST-RAD54B-PM (S4-E33) and DMC1, either wild type or with the indicated amino acid substitutions. **(E)** Apparent affinities for the ssDNA-DMC1 filaments, either WT or mutated, measured by BLI for both BRCA2-P and RAD54B-P (see also Table S2 and Figure S5D). **(F)** Electromobility Shift assay (EMSA) of DMC1 filaments, in absence or in presence of increasing concentrations of BRCA2-PL or RAD54B-PL (0.25, 0.5 and 1 µM). The supershift shows that the peptide binds to the ssDNA-DMC1 filaments. **(G)** Filament lengths measured for DMC1 WT and mutated in the absence (grey) and presence of BRCA2-PL (blue) and RAD54B-PL (pink).

We searched for the molecular basis of this stabilization effect. Therefore, we made two different but not exclusive hypotheses: (1) stabilization comes from specific contacts that mediate preferential binding of the peptides to DMC1 in its filament-assembled conformation; (2) stabilization comes from transient contacts between the positively charged residues of the peptides and negative patches distributed on different DMC1 protomers. To challenge these hypotheses and identify DMC1 residues essential for stabilization by P-motif peptides, we mutated negatively charged DMC1 residues (**Figure 4C**).

Within mutations of DMC1 negatively charged residues, we previously identified E213K as disrupting most the binding to BRCA2 G2379-Q2433 (**30**). Here, we performed GST-pulldown analyses between DMC1 mutants and RAD54B-PM. We found again that, among all designed mutants, only E213K strongly disrupts binding (**Figure 4D; Figure S5A**). In our cryo-EM structures, E213 is in front of a positively charged residue in BRCA2 (K2404) but not in RAD54B (P18) (**Figure 4C**). Thus, our data suggest that mutation E213K does not disrupt a specific salt bridge between DMC1 and one of the peptides. To evaluate more accurately the impact of the DMC1 mutations on the filament binding to the peptides, we purified all the mutants, verified that they were correctly folded (**Figures S5B, C**), and ran BLI experiments with the peptides BRCA2-P and RAD54B-P (**Figure 4E; Figure S5D; Table S2**). Even though DMC1 E213K could be loaded onto ssDNA (**Figures S5D, E**), we could not measure any affinity value for the interaction between these filaments and either of the peptides, in agreement with the GST-pulldown experiments. The BLI data also revealed that the mutations D175K and D180K mildly reduced the affinity of the filaments for both peptides: D180K decreased the affinity by 4-fold and D175K by 2-fold, whereas our control mutation F89A (F89 being at the interface between 2 protomers) did not significantly affect these interactions. Unexpectedly, for all mutants, we did systematically observe the same affinity decrease for BRCA2-P and RAD54B-P, suggesting that D175K, D180K and E213K either modify the overall long-range electrostatic interactions with the peptides or affect the structure of the DMC1 binding site.

Finally, we tested if the mutations modified the filament stabilization by BRCA2-PL and RAD54-PL. In conditions in which filaments are assembled and bind to the peptides, as confirmed by negative-staining EM (**Figure S5E**) and electrophoretic mobility shift assays (EMSA) (**Figure 4F**), we observed that F89A did not impact the capacity of the filaments to be stabilized by BRCA2-PL, but all 3 other mutations (D175K, D180K, E213K) completely abolished this stabilization effect (**Figure 4G**). Similarly, mutation E213K disrupted the stabilizing effect of RAD54B-PL on ssDNA-DMC1 filaments. Thus, mutating the DMC1 negatively charged residues located at and around the P-site decreased binding to the BRCA2-P and RAD54B-P peptides and eliminated their stabilizing impact.

### The N-terminal region of DMC1 contributes to the affinity of the BRCA2 and RAD54B peptides for the DMC1 nucleoprotein filaments

The BRCA2-PL and RAD54B-PL peptides bind to a P-site located on the ATPase domain of a DMC1 protomer, but also adjacent to its N-terminal disordered and folded regions (**Figures 1A, 3A**). To explore the role of the N-terminal region of DMC1 in the stabilization of the presynaptic filaments by BRCA2 and RAD54B, we tested experimentally the impact of deleting the DMC1 disordered and folded domains. Only DMC1 lacking the first 18 residues (ΔN18), but not DMC1 lacking its entire N-terminal region (ΔN81), still assembles into filaments in our conditions, as shown by EMSA (**Figure S6A**). Moreover, deletion of the DMC1 18 first residues reduces the apparent affinity of the filament for BRCA2-P by 2-fold, and for RAD54B-P by more than 3-fold (**Figure 5A; Figure S6B; Table S1**). Finally, the DMC1 ΔN18 filaments lost their capacity to be stabilized by both BRCA2-PL and RAD54B-PL (**Figure 5B**).

**Figure 5.**
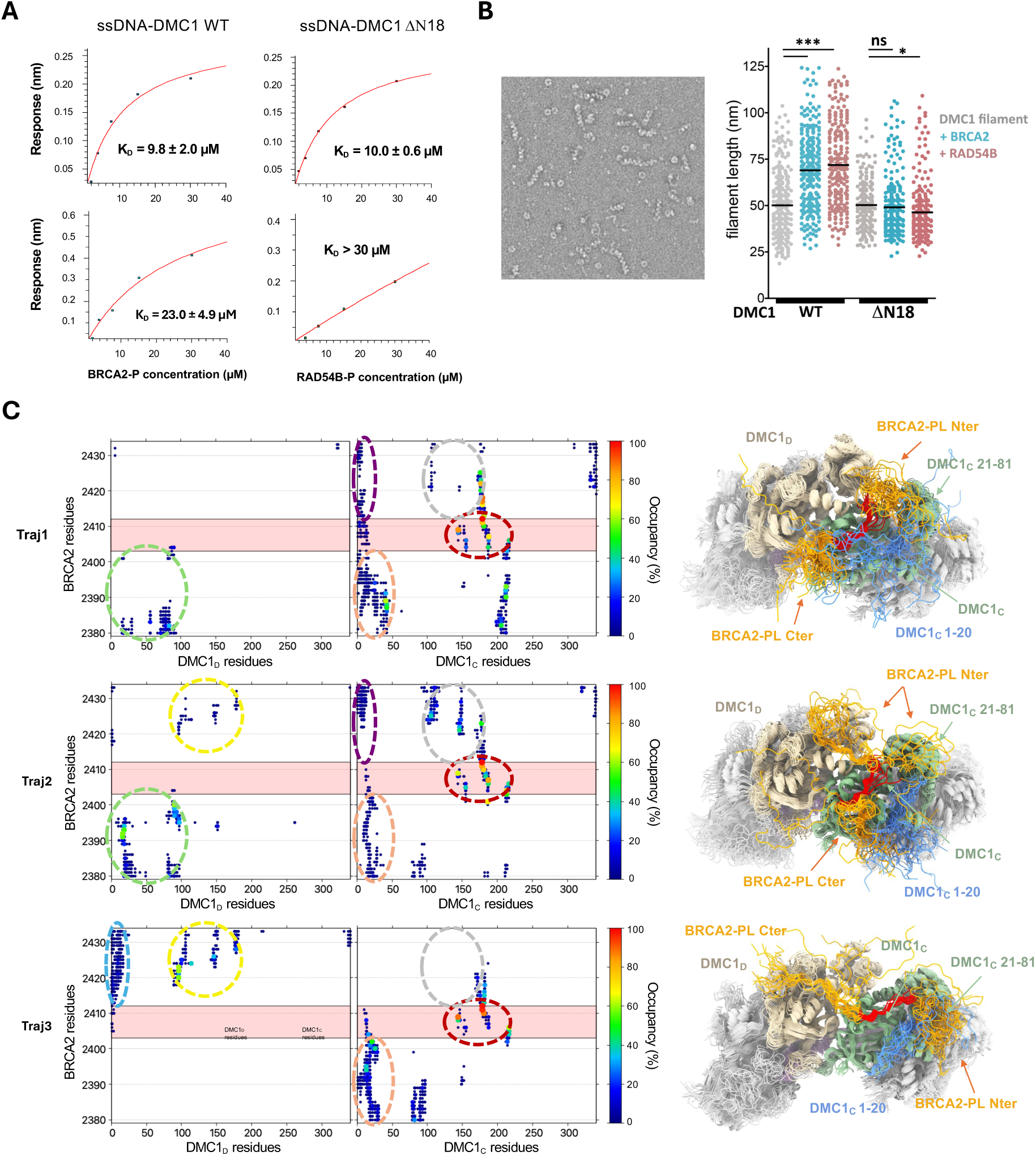
The negatively charged N-terminal disordered region of DMC1 contributes to binding and stabilization of the ssDNA-DMC1 filaments. **(A)** Apparent affinities for the ssDNA-DMC1 filaments, either WT or DN18, measured by BLI for both BRCA2-P and RAD54B-P (see Table S2 and Figure S6B for replicates). **(B)** Comparison of the ssDNA-DMC1 filament lengths, as measured by negative-staining EM, for DMC1 WT or DN18 in the absence and presence of the peptides. **(C)** Contact maps and representative structures showing the proximities between BRCA2-PL and the bound (DMC1C) or adjacent (DMC1D) recombinases along the 3 MD trajectories calculated for the ssDNA-DMC1-BRCA2-PL filament (see Figs. S8-10). On the left, the contacts corresponding to the cryoEM structure are surrounded by red dashed lines, whereas the contacts systematically detected between the N-terminus of the peptide and the N-terminal of DMC1 are surrounded by orange dashed lines. Additional contacts are identified using dashed circles of different colors. On the right, representative structures of the 20 clusters extracted for each trajectory are superimposed. DMC1 monomers A, B, E, and F are colored light grey, monomer D is colored light brown, monomer C is colored green, except for residues 1–20 that are colored blue. Residues of the BRCA2 peptide are colored in orange except for residues 2403 to 2412 (observed in our cryoEM structure) that are colored red. For clarity, residues 1–30 of monomers A, B, D, E, and F are omitted.

To decipher the role of the N-terminal region of DMC1 in peptide binding and filament stabilization, we turned to molecular modeling. First, using AlphaFold3 (AF3), three-dimensional models of complexes between a DMC1 dimer and several BRCA2 and RAD54B peptides were calculated. All these models were consistent with our cryo-EM structures: DMC1 adopts a filament-assembled structure, and the core sequences of the peptides are docked at the DMC1 P-site, as illustrated in **Figure S7**. However, the positions of the residues that were not observed by cryo-EM at the N-terminus of DMC1 and on both sides of the core sequences in the peptides were predicted with low confidence, suggesting that these residues adopt multiple conformations. Proximities were still observed between the DMC1 N-terminal region and the peptides (**Figure S7**). This encouraged us to further investigate the interactions involving the DMC1 N-terminal region using molecular dynamics (MD) simulation. We modeled an 18-nt ssDNA covered with 6 DMC1 (DMC1_A_ to DMC1_F_) and positioned a BRCA2-PL peptide on DMC1_C,_ its core sequence being bound as observed by cryo-EM (**Figure S8A**). We then performed 3 MD trajectories of this DMC1 presynaptic filament in an explicit solvent environment. For each trajectory, root-mean-square fluctuations (RMSF) were calculated around the mean structure to characterize the dynamics of the DMC1 protomers and BRCA2 peptides. The RMSF profiles of protomers B, C, and D are very similar. Residues M1 to S19 display very large RMSF values in all three trajectories, which is consistent with the lack of electron density for these residues in the cryo-EM maps (**Figure S8B**). The BRCA2 residues identified by cryo-EM have restricted fluctuations, whereas the distal parts of the peptide display large RMSF values (**Figure S8C**). To analyze the BRCA2 interactions during the simulations, we constructed contact maps between BRCA2 and the DMC1 protomers B, C and D (**Figure 5C, left panels; Figure S8D**). Analysis of these maps revealed that (i) interactions involving the BRCA2 residues detected by cryo-EM and the DMC1 P-site display the highest occupancies in the 3 trajectories (green to red dots in red circles in **Figure 5C**), (ii) the N-terminus of BRCA2-PL interacts with the N-terminal region of DMC1_C_ in all trajectories, but these contacts have lower occupancies (blue to green dots in orange circles in **Figure 5C**), (iii) different regions of BRCA2-PL and DMC1_D_ interact in each of the 3 trajectories, and no contacts are detected between BRCA2-PL and the other DMC1 protomers (**Figure 5C; Figure S8D**). Finally, we performed a cluster analysis of the 3 trajectories and extracted representative structures of the conformations sampled during the simulations (**Figure 5C, right panels; Figure S8E**). Consistently with the analysis of the contact map, we observed that the N-terminus of the BRCA2 peptide transiently interacts with the N-terminal folded domain of DMC1_C_ in all trajectories (orange circles in the left panels), and sometimes with the N-terminal domain and the linker of DMC1_D_ (traj. 1 & 2; green circles), whereas the C-terminus of the peptide interacts with the ATPase domain of DMC1_C_ (grey circles), and sometimes with the disordered N-terminal region of DMC1_C_ (traj. 1 & 2; violet circles) or DMC1_D_ (traj. 3; cyan circles) as well as the ATPase domain of DMC1_D_ (traj. 2 & 3; yellow circles). To illustrate the role of the BRCA2-PL positively charged residues in these transient interactions, we monitored specific proximities involving BRCA2 K2404 and K2413 during the trajectories (**Figures S9 & S10**). We found that K2404 may transiently form a salt bridge with the N-terminal residue E16 (traj. 2 & 3) and the ATPase domain residues K155, D186 (traj. 1 & 2) and E213 (traj. 2) of DMC1_C_ as well as with E90 in the linker region of DMC1_D_ (traj. 1). Similarly, K2413 may form a salt bridge with the ATPase domain residues D175 (traj. 3), D180 and D182 (all traj.) of DMC1_C_ as well as with D4 in the N-terminal disordered region of DMC1_D_ (traj. 1). Altogether, our MD simulations revealed a dense network of interactions between the distal regions of the peptide and the N-terminal and ATPase domains of DMC1_C_, as well as the N-terminal region of DMC1_D_.

## Discussion

The ssDNA-DMC1 filament is an essential actor of meiotic recombination. DMC1 is expressed from the very beginning of meiosis, in prophase I, when programmed DNA double-strand breaks are introduced. The ssDNA-DMC1 filament is required for homologous synapsis of chromosomes (55, 56). Its assembly and disassembly are strongly regulated: if it is too unstable, the probability of finding homologous sequences is reduced; conversely, an excessively stable filament is not efficient at forming the D-loop and promoting strand exchange. Accessory proteins help stabilize the filament while maintaining its functional dynamics (57). BRCA2, Hop2-Mnd1 and Swi5-Sfr1 regulate DMC1 filaments, although no detailed structural mechanism was reported to support their functions (26, 28, 58, 59). Here we focused on the interaction between BRCA2, RAD54B, and the meiotic ssDNA-DMC1 filament. We determined high-resolution cryo-EM structures of BRCA2 and RAD54B selected fragments bound to a ssDNA-DMC1 filament and asked how these peptides stabilize the meiotic recombinogenic filament.

### High-resolution cryo-EM structures of BRCA2-PL and RAD54B-PL bound to ssDNA-DMC1 reveal a very similar core binding motif but do not recapitulate all the contacts contributing to the peptide-filament interactions

By solving the 3D structures of a 43-aa BRCA2 peptide and a 29-aa RAD54B peptide bound to ssDNA-DMC1 at a resolution of 1.9-2.0 Å, we identified the interface between the peptides and the filament. We observed that it was mostly hydrophobic, with only two conserved hydrogen bonds between the backbone atoms of the peptide C-termini and DMC1 residues N178 and D180 (**Figure 3D**). Unexpectedly, no high occupancy salt-bridge interaction was detected between the positively charged peptides and the negatively charged DMC1 P-site in the cryo-EM structures as well as in the MD trajectories. Still, mutating K2404 or K2413 in BRCA2-P and K16, K17 or K19 in RAD54B-P significantly decreased binding to the DMC1 filament ((30); **Figure 2E, F**). These positively charged residues are conserved, but they are not located at the same positions in the two peptides: they could mediate specific interactions with different residues of DMC1 (**Figure 4C**). However, mutating D175, D180 and E213 located in front of these positively charged residues in DMC1 caused the same affinity decrease for both BRCA2-P and RAD54B-P peptides (**Figure 4E**), showing that, consistently with the cryo-EM and MD analyses, the mutated residues do not mediate a specific interaction with one of the peptides. Based on these data, we propose that electrostatic interactions between the BRCA2-P and RAD54B-P peptides and the DMC1 protomers are multiple and transient. They could still contribute to the binding specificity towards DMC1 versus RAD51 filaments, as residues at the periphery of the F-x-P-P docking site (ex: D180, E213) are not conserved between the two recombinases (**Figure 2A**).

### The BRCA2-PL and RAD54B-PL peptides transiently interacts with the N-terminal disordered and folded regions of DMC1 of two adjacent protomers

By further analyzing the interaction between the P-motif containing peptides and DMC1 filaments using AF3, we hypothesized that the peptides could also transiently interact with the negatively charged N-terminal region of DMC1. Consistently, we found that deleting the disordered 18 first residues of DMC1, which contains 8 negatively charged residues conserved in vertebrates, decreases the affinity between the DMC1 filament and the peptides (**Figure 5A**), and abolishes filament stabilization by BRCA2-PL and RAD54B-PL (**Figure 5B**). The role of the N-terminal folded domain of DMC1, which is mostly positively charged but also contains few negatively charged patches including D23 & D25, is more difficult to test experimentally as its deletion prevents filament formation. MD simulations showed that not only the disordered N-terminal region of DMC1_C_, but also the folded N-terminal domains of DMC1_C_ and DMC1_D_, might transiently contact the disordered N-terminal residues of the BRCA2-PL peptide (**Figure 5C**). This analysis suggests that electrostatic interactions between the entire N-terminal region of DMC1 and the peptides may contribute to stabilize the ssDNA-DMC1 filaments by not only increasing the affinity between filaments and peptides but also transiently bridging the N-terminal folded domains of two adjacent DMC1 protomers.

### The filament-specific conformation of the N-terminal region of DMC1 might contribute to filament stabilization by BRCA2-PL and RAD54B-PL

To explain why the BRCA2 PhePP core motif R2401-S2414 specifically recognizes oligomeric DMC1 and further stimulates filament formation, Dunce and Davies proposed that peptide binding may allosterically favor the ssDNA-bound DMC1 conformation (31). Similarly, it was suggested that TR2 could allosterically reshape the RAD51 conformation to favor dsDNA binding (60). BRCA2-PL and RAD54B-P bind to the ssDNA-DMC1 filament without inducing any detectable changes in the three-dimensional structure of DMC1. However, several regions involved in the interaction are not observed by cryo-EM. Our BLI and MD results suggest that the N-terminal residues of the peptides interact with the N-terminal disordered and folded sequences of DMC1. The DMC1 N-terminal region adopts different conformations as a function of the structural context. In the DMC1 octamer structure, it is not detected because its position relative to the ATPase domain is variable (18). Upon binding of the DMC1 octamer to BRCA2-P, its position is partially stabilized: the DMC1 region from Q28 to L80 is detected, even if with high B-factors, and is docked against a negatively charged patch in the ATPase domain of the adjacent protomer (30). When DMC1 is assembled into a filament, its N-terminal folded domain adopts a different and more stable position: it is docked against a second negatively charged patch in its own ATPase domain and a third negatively charged patch on the adjacent ATPase domain. This filament-specific position of the N-terminal region of DMC1 could favor transient interactions between filaments and peptides, thus, in turn, contributing to the stabilization effect of the peptides.

## Conclusion

Here, we showed that both BRCA2-PL and RAD54B-PL can stably anchor onto the ssDNA-DMC1 filament. The interfaces between the peptides and the filament share a central hydrophobic core in which the motif F-[I/V]-P-P binds to the P-site of a single DMC1 protomer, and additional transient electrostatic contacts between positively charged residues of P-motifs and negatively charged residues of DMC1. We propose that the hydrophobic anchor allows N- and C-terminal distal regions of the peptides to establish transient electrostatic interactions with the N-terminal and ATPase domains of DMC1. This highlights that the definition of the P-motif includes not only the F-x-P-P sequence, but also essential lysine and/or arginine residues, whose presence should always be taken into account when screening for P-motifs in unstructured regions of recombinase-interacting partners.

*In vitro*, BRCA2-PL and RAD54B-PL stabilize the ssDNA-DMC1 filament. Our cryo-EM analysis did not identify persistent contacts between the peptides and adjacent DMC1 protomers in the filament, as reported in the case of BRCA2 (TR2) and RAD51AP1 interacting with ssDNA-RAD51 (61). However, mutating DMC1 negatively charged residues or deleting the DMC1 N-terminal disordered and negatively charged region abolished the stabilizing effect of the peptides. We propose that the positively charged peptides transiently contact the N-terminal domain and the linker of the adjacent protomer. Additionally, RAD54B interacts through another region, located between S26 and R225, with DMC1 (38). Thus, RAD54B might also bridge two protomers through two different DMC1 binding sites. Finally, it was very recently reported that RAD54B exhibits three different sites binding to ssDNA-RAD51 filaments, the anchoring site comprising the three first residues of RAD54B, consistently with our pull-down data (62). The authors suggested that RAD54B takes advantage of its three binding sites to bridge two protomers within the ssDNA-RAD51 filaments. All these interactions could contribute not only to filament stabilization but also to increasing the specificity of the binding to the nucleoprotein filament. Further studies should identify the steps at which BRCA2, RAD54B, and DMC1 functions are coupled to yield an active ssDNA-DMC1 filament in the highly organized meiotic chromatin context.

## Acknowledgments

We warmly thank the cryo-EM facility of I2BC, and especially Laura Pieri and Stéphane Bressanelli, for their help during data acquisition and processing, and the Macromolecular Interaction Platform of I2BC, and especially Magali Nicaise-Aumont and Magali Noiray, for their help on the BLI instrument. We acknowledge SOLEIL for provision of cryo-EM facilities on the POLARIS TITAN KRIOS. We thank Chloé Quignot for implementing the AlphaFold-Multimer at I2BC.

## Author contributions

S.R.F., A.M. and M.L.H. purified and characterized the proteins. A.Z. did bioinformatic analyses, found and performed initial characterization of the DMC1-binding motif of RAD54B. S.R.F. performed the GST-pulldown analyses. S.M. performed the BLI and ITC analyses. P.D., S.B., M.O. and P.L. prepared the cryo-EM grids and acquired the cryo-EM data. S.B., S.Z.J. and P.L. processed the cryo-EM data. A.M., S.M. and P.D. acquired the EMSA, D-loop and negative-staining EM data. P.C. did the MD simulation analysis. P.D., S.M., P.C., A.Z. and S.Z.J. designed the experiments and analyzed the data. S.Z.J and R.K. acquired funding and supervised the work. S.Z.J. wrote the original draft of the manuscript. All authors contributed to the writing, reviewed, and edited the manuscript.

## Data availability

Cryo-EM maps and structural models have been deposited in PDB (32RS, 32TK, 32TL) and EMDB (EMD-59127, EMD-59147, EMD-59148).

