## Supplementary information for "BRCA2 and RAD54B FxPP motifs Bind DMC1 Filaments through Persistent and Transient Interfaces"

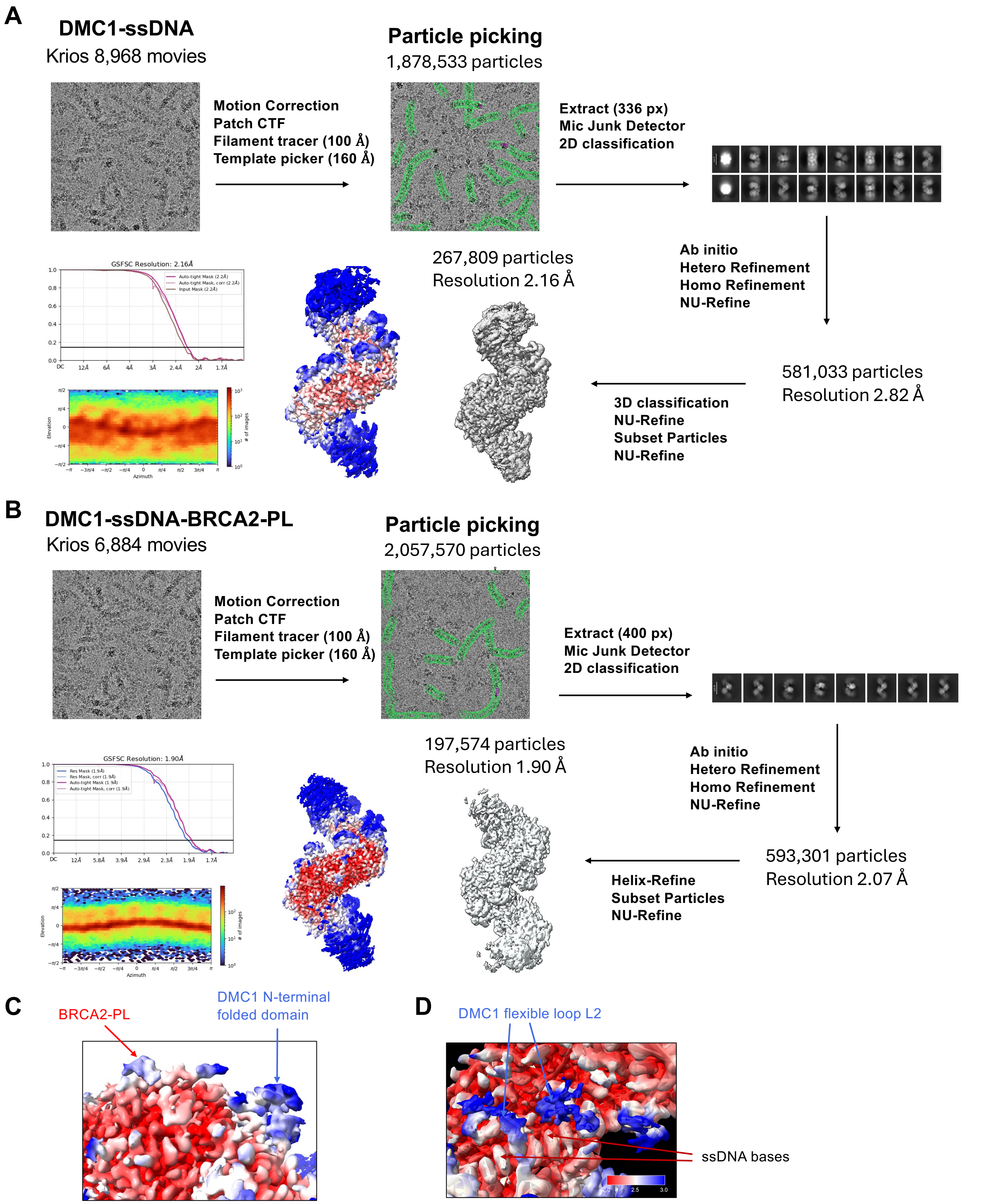


**Figure S1. Related to Figure 1. Determination of the cryoEM structures of ssDNA-DMC1 and ssDNA-DMC1-BRCA2-PL. (A)** Data processing flowchart of the ssDNA-DMC1 filaments (AMP-PNP and $Ca^{2+}$) without any bound peptide. The colored EM map shows the local resolution, from 2 (red) to 3 (blue) Å. **(B)** Data processing flowchart of the $Ca^{2+}$-ATP ssDNA-DMC1 filaments in the presence of BRCA2-PL G2379-Q2421. The local resolution is illustrated as in (A). **(C)** Zoom view from (B) focused on the the BRCA2 peptide. **(D)** Zoom view from (B) focused on the DMC1 DNA binding loop L2 and ssDNA.

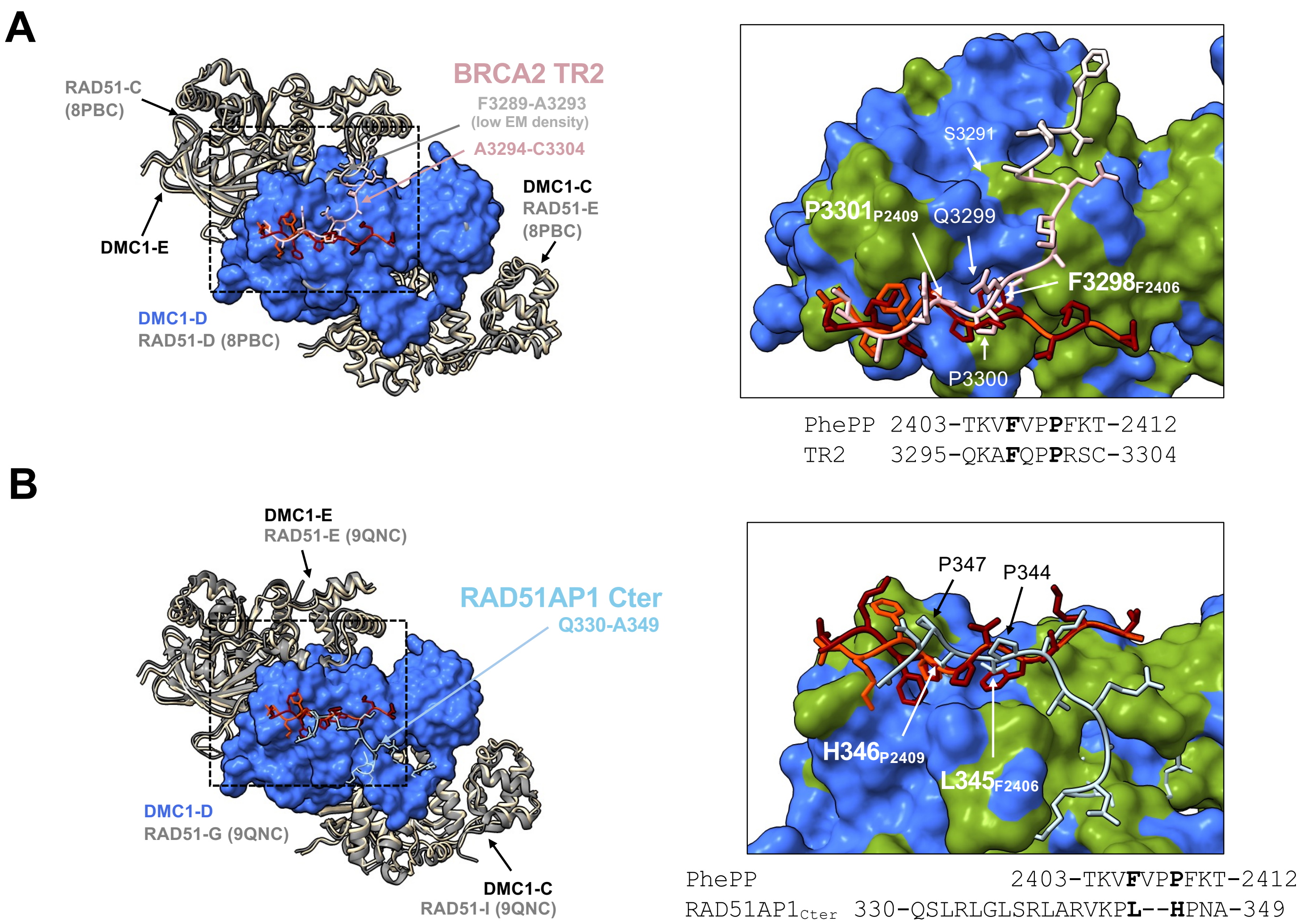


**Figure S2. Related to Figure 1. Comparison between the DMC1 and RAD51 P-sites. (A)** Superimposition of the BRCA2 PhePP structures bound to the ssDNA-DMC1 filament and the BRCA2 TR2 structure bound to the ssDNA-RAD51 filament (8PBC; resolution 2.6 Å; ATP and $Ca^{2+}$): BRCA2 F2406 and F2409 superimpose onto TR2 F3298 and P3301, respectively. **(B)** Superimposition of the PhePP structures bound to the ssDNA-DMC1 filament and the RAD51AP1 C-terminus bound to the ssDNA-RAD51 filament (9QNC; resolution 3.0 Å; ATP and $Mg^{2+}$): the BRCA2 residues F2406 and P2409 bind into the same recombinase cavity as the RAD51AP1 residues L345 and H346. The BRCA2 and RAD51AP1 peptides are shown as cartoons, with their side chains in sticks. In the zoom panels, the DMC1 surface is colored in royal blue, except for residues not conserved in RAD51 that are in dark green.

**A**

```
RAD54B-P    (12-29)              GNSFKKPKFIPPGRSNPG
RAD54B-PM   (4-33)        SAAPSQLQGNSFKKPKFIPPGRSNPGLNEE
RAD54B-PL   (1-29)     MRRSAAPSQLQGNSFKKPKFIPPGRSNPG
RAD54B-PXL  (1-33)     MRRSAAPSQLQGNSFKKPKFIPPGRSNPGLNEE
```

**B**

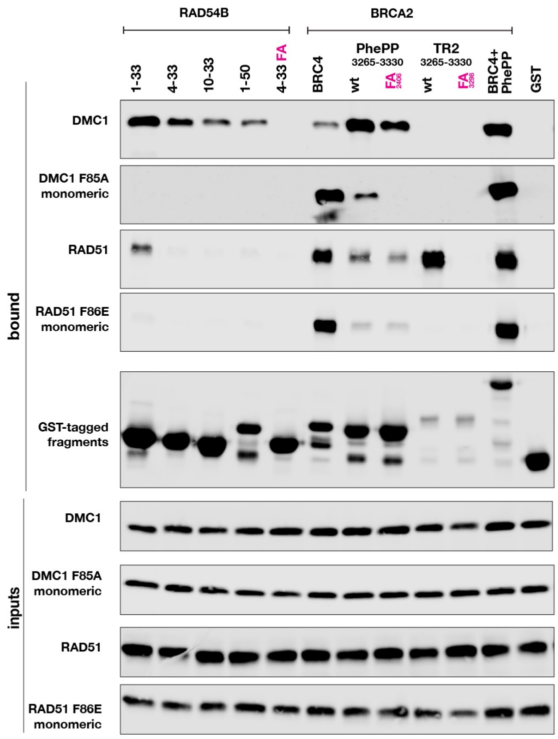


**C**

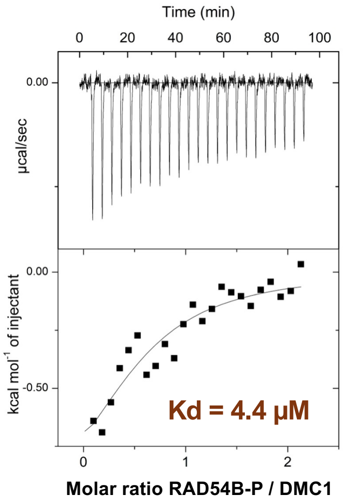


**D**

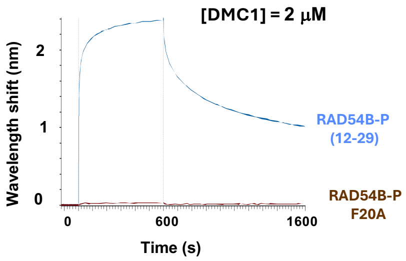


**Figure S3. Related to Figure 2. Characterisation of the interaction between the RAD54B N-terminal region and the DMC1 octamer. (A)** Sequences of RAD54B peptides tested in this study. The conserved residues, as defined in Figure 2B, are in bold. The residues mutated in this study are in blue and red. **(B)** Replicate of the experiment in Figure 2C. **(C)** ITC experiment showing that RAD54B binds to DMC1 octamers with a micromolar affinity. A binding affinity of 4.4 +/- 6.6 µM was measured with RAD54B-P. **(D)** BLI experiments testing the binding of immobilized RAD54B-P peptides, wild-type and mutated F20A, to octameric DMC1.

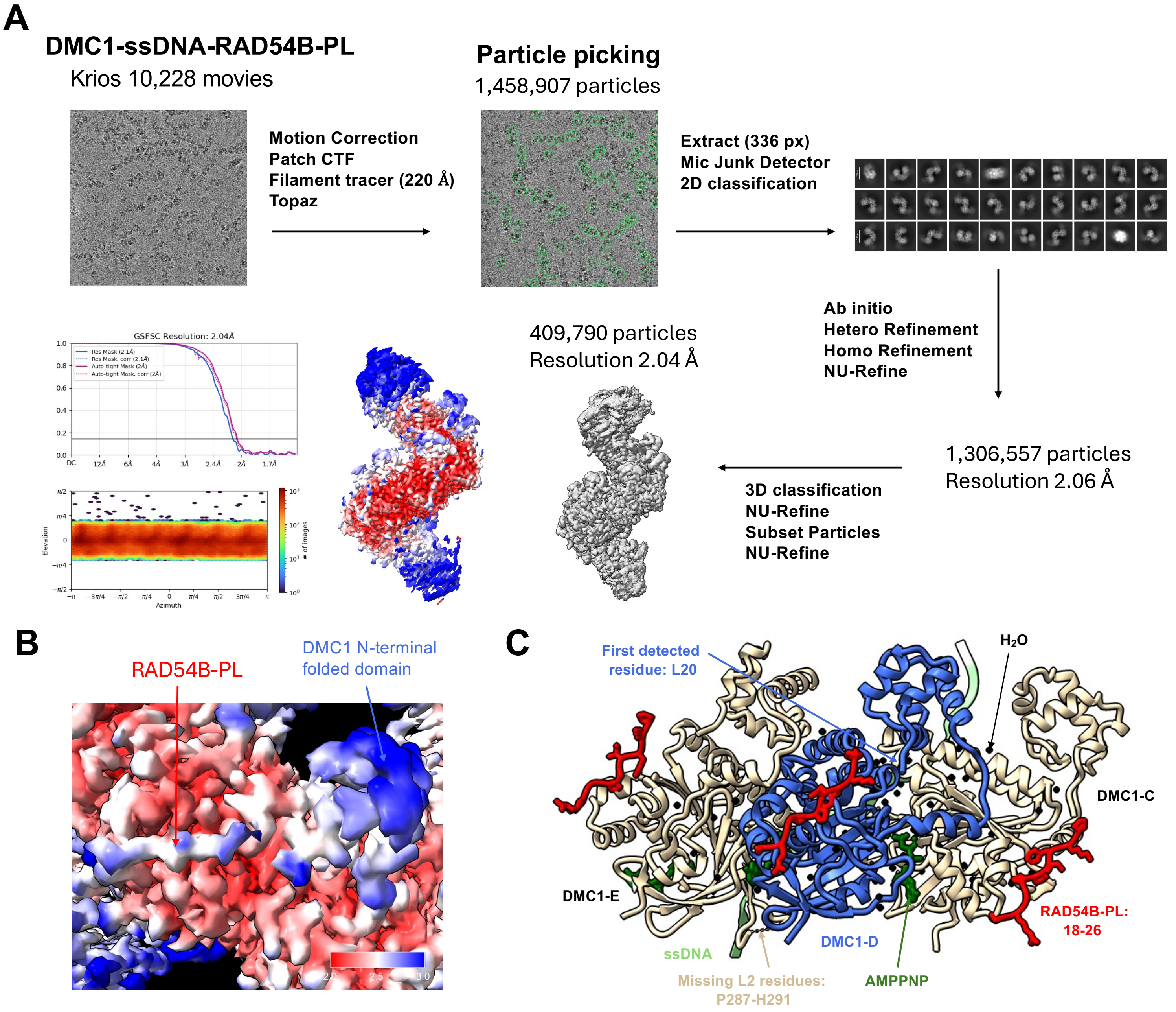


**Figure S4. Related to Figure 3. Determination of the high resolution cryoEM structure of ssDNA-DMC1 in the presence of RAD54B-PL. (A)** Data processing flowchart of the ssDNA-DMC1 filaments (AMP-PNP and $Ca^{2+}$) interacting with RAD54B-PL M1-G29. The local resolution of the map is indicated using a color scale going from 2 (red) to 3 (blue) Å. **(B)** Zoom view from (A) focused on the the RAD54B-PL peptide. **(C)** View of the ssDNA-DMC1-RAD54B-PL cryoEM structure, showing only three DMC1 protomers.

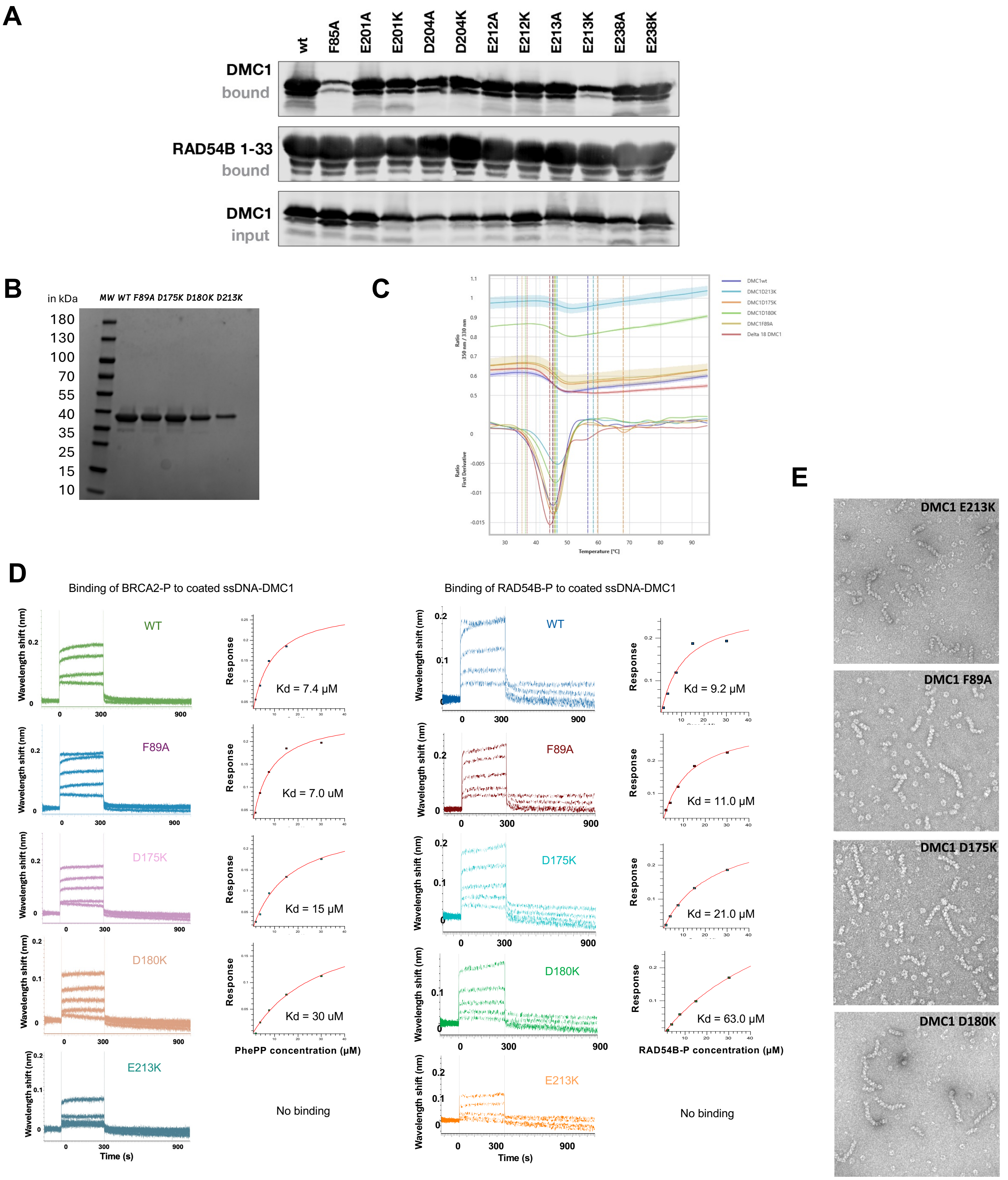


**Figure S5. Related to Figure 4. Analysis of the interaction between the mutated ssDNA-DMC1 filaments and the P-motif peptides. (A)** GST-pulldown experiment as in Figure 4D, but with RAD54B-PXL (M1-E33). **(B)** SDS-PAGE gel of the purified DMC1 proteins, wild-type and mutated. **(C)** Thermal stability of DMC1 variants determined by nanoDSF. Thermal unfolding profiles of DMC1 WT, F89A, D175K, D180K, E213K and Δ18 were recorded using a Prometheus Panta instrument (Nanotemper). The curves represent the first derivative of the intrinsic fluorescence ratio (F350/F330) as a function of temperature. The minima correspond to the apparent melting temperatures (Tm), and the dashed vertical lines indicate the Tm values calculated for each protein variant. **(D)** Measurement of the affinities of BRCA2-P and RAD54B-P for the WT and mutated filaments by BLI. After loading similar amounts of all the DMC1 variants onto ssDNA, the BRCA2-P or RAD54B-P peptide was progressively added to the filaments. The kinetics of association and dissociation of the interaction was recorded for each peptide concentration, in order to calculate a Kd value in the steady-state mode. **(E)** Negative-staining EM images of the ssDNA-DMC1 filaments formed by mutated DMC1 recombinases.

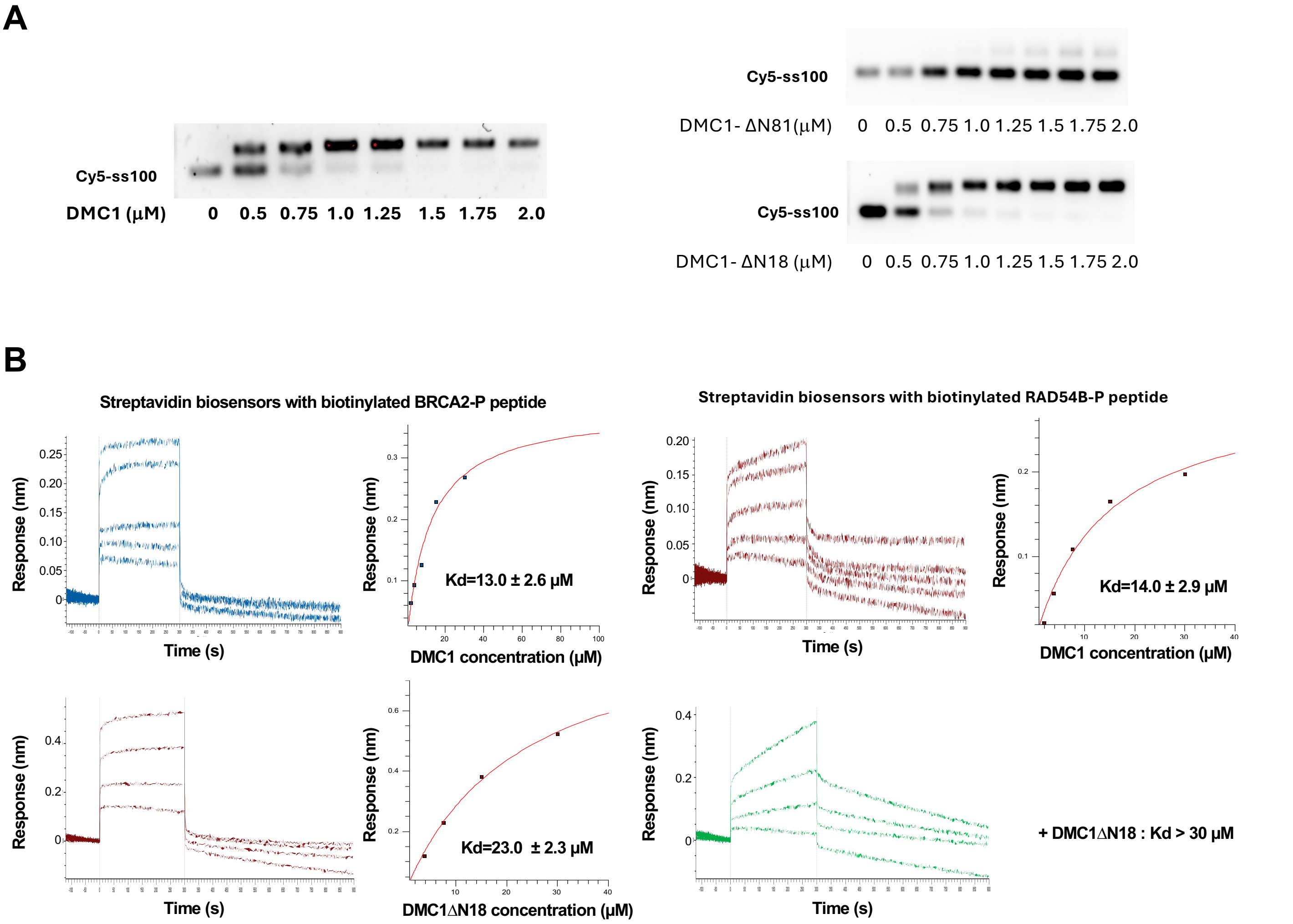


**Figure S6. Related to Figure 5. Analysis of the interaction between the filaments assembled from truncated DMC1 proteins and the P-motif peptides. (A)** Electromobility Shift assay (EMSA) of a Cy5-100-nt ssDNA, in presence of increasing concentrations of DMC1, either WT (left) or truncated (right). The supershift shows that it was possible to assemble nucleoprotein filaments with DMC1 WT and ΔN18, but not with DMC1 ΔN81. **(B)** Replicates of the BLI experiments performed to measure the apparent affinities of BRCA2-P and RAD54B-P for the ssDNA-DMC1 filaments, assembled from either DMC1 WT or DMC1 ΔN18.

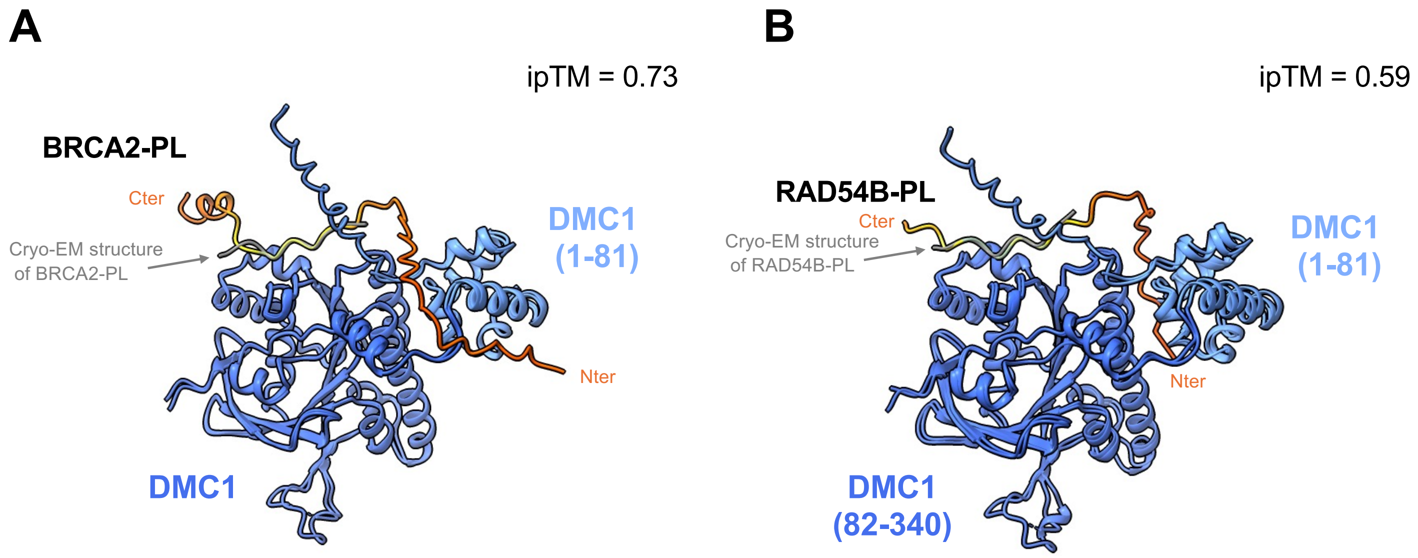


**Figure S7. AlphaFold3 models of the interaction between P-motif containing peptides and presynaptic recombinase filaments. (A) The 43-aa BRCA2-PL peptide was docked onto a dimer of DMC1 bound to ssDNA. (B) The 29-aa RAD54B-PL peptide was docked onto a dimer of DMC1 bound to ssDNA.** A representative model (within the 5 calculated models) of each AF complex is superimposed onto the cryo-EM structure of the corresponding complex. Only one recombinase and one peptide are displayed for clarity. The recombinase N-terminal domains (1-81) are in cornflower blue and their ATPase domains (82-340) are in royal blue. The cryo-EM peptides are in grey and the AF peptides are colored as a function of their pLDDT.

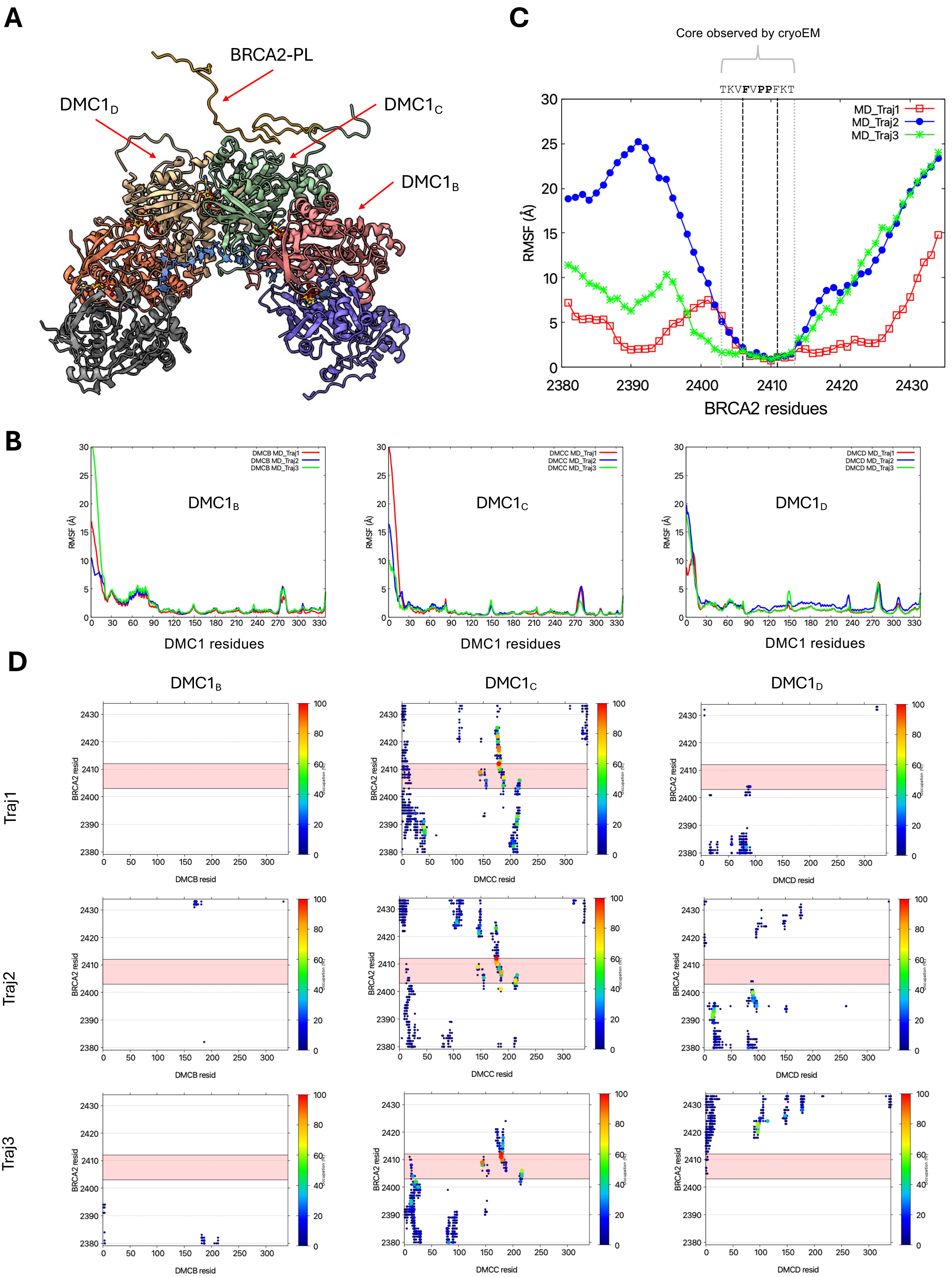
A
BRCA2-PL
DMC1$_D$
DMC1$_C$
DMC1$_B$
C
Core observed by cryoEM
TKVFVPPFKT
MD_Traj1
MD_Traj2
MD_Traj3
RMSF (Å)
BRCA2 residues
B
DMCB MD_Traj1
DMCB MD_Traj2
DMCB MD_Traj3
DMCC MD_Traj1
DMCC MD_Traj2
DMCC MD_Traj3
DMCD MD_Traj1
DMCD MD_Traj2
DMCD MD_Traj3
RMSF (Å)
DMC1 residues
D
Traj1
Traj2
Traj3
BRCA2 resid
DMCB resid
DMCC resid
DMCD resid

**E**

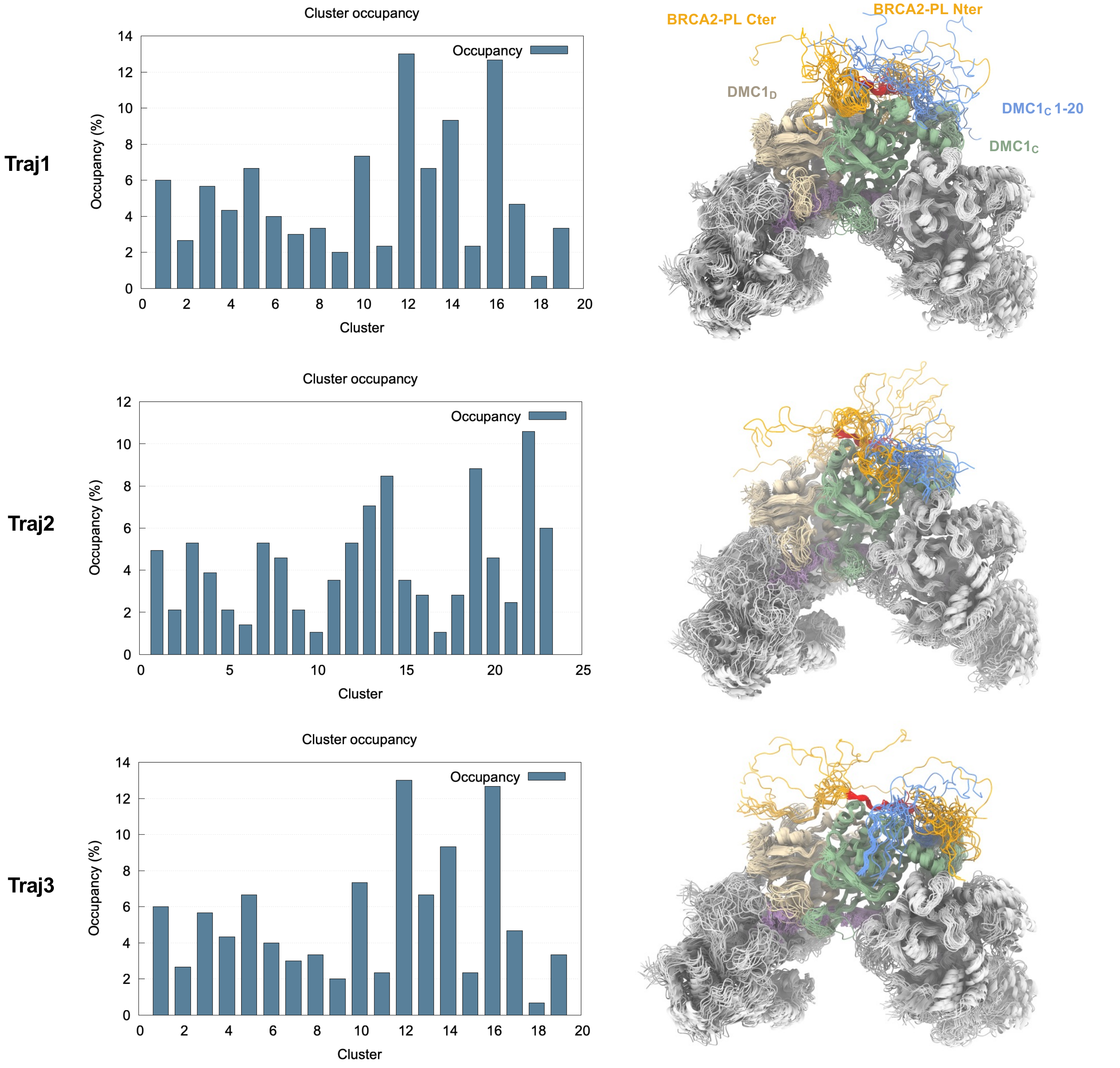


**Figure S8. Extended data on the MD simulations of Figure 5. (A)** Initial ssDNA-DMC1-BRCA2-PL structure for MD simulation. **(B)** Root Mean Square Fluctuation (RMSF) of the coordinates of the Ca atoms of three DMC1 protomers, always after superimposition on the central and BRCA2-bound DMC1$_C$. **(C)** RMSF of the coordinates of the Ca atoms of the BRCA2 peptide along the three MD trajectories calculated for the ssDNA-DMC1-BRCA2-PL filament. The reported values are calculated after superimposition on the monomer DMC1$_C$ that interacts with the BRCA2 peptide. **(D)** Full contact map between PhePP and the bound (DMC1$_C$) or adjacent (DMC1$_B$, DMC1$_D$) recombinases deduced from the analysis of 3 independent molecular dynamics simulation trajectories of 300 ns. **(E)** Cluster analysis of molecular dynamics simulations of the ssDNA-DMC1-BRCA2-PL complex**.** The populations of the clusters identified in trajectories 1, 2, and 3 are shown in the left panels. Side views and top views of the BRCA2-binding site are shown in the right panels: for each trajectory, 20 representative cluster conformations from trajectories 1, 2, and 3 are superimposed onto DMC1$_C$. In these views, DMC1 monomers and the BRCA2 peptide are shown in cartoon representation. DMC1 monomers A, B, E, and F are colored light grey, monomer D is colored light brown, monomer C is colored green, except for residues 1–20 that are colored blue. Residues of the BRCA2 peptide are colored in orange; except for residues 2403 to 2412 that are colored red. For clarity, residues 1–30 of monomers A, B, D, E, and F are omitted.

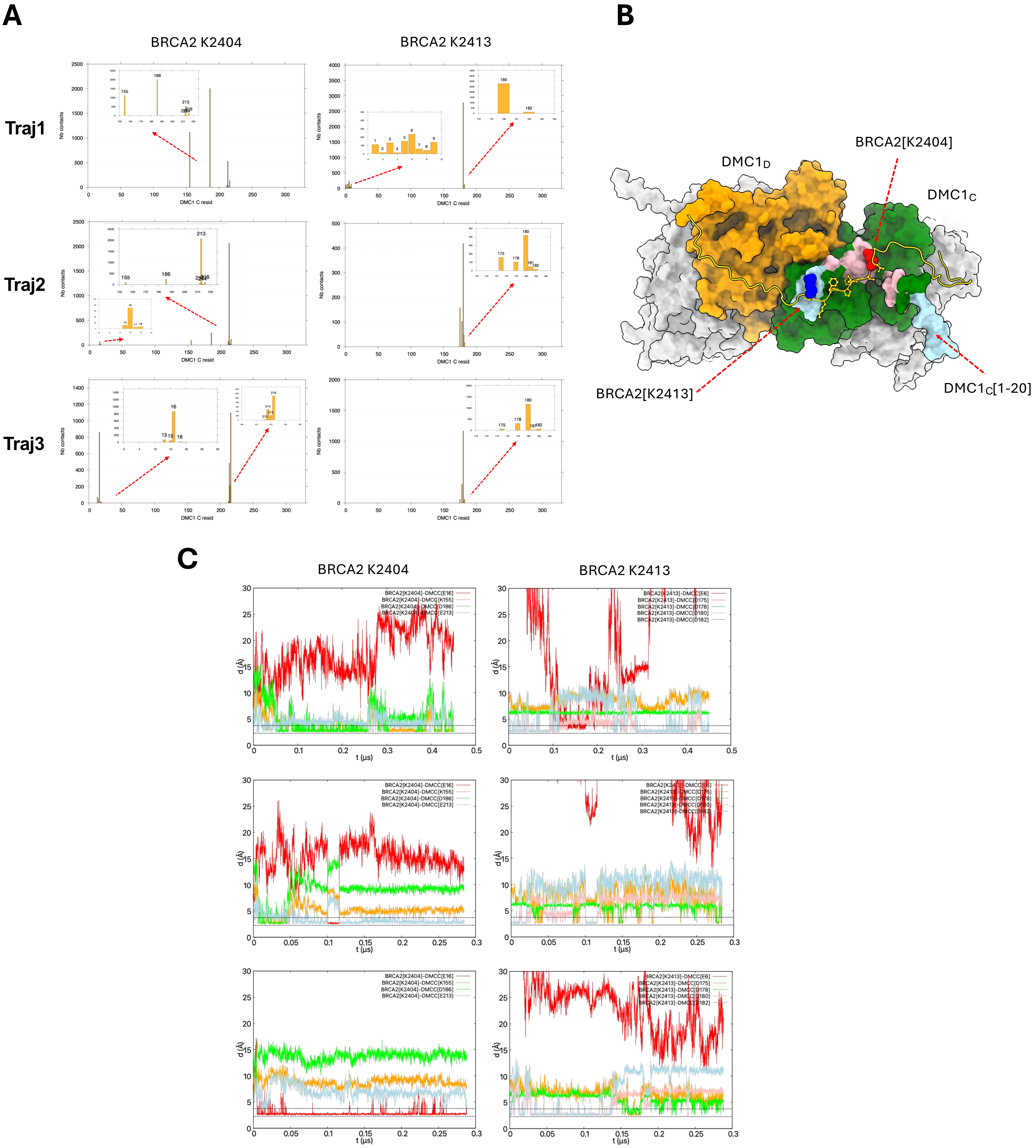


**Figure S9. Extended data on the MD simulations of Figure 5. Contact analysis of BRCA2-PL K2404 and K2413 in molecular dynamics simulations of the ssDNA–DMC1-BRCA2-PL complex: interactions with DMC1$_C$. (A)** Residues of the DMC1$_C$ protomer contacting BRCA2 residue K2404 or K2413 are shown together with the number of frames in which each contact (d<4.0Å) was detected. Frames were sampled every 0.1 ns throughout the simulations. **(B)** Contacts of panel (A) are mapped onto the starting structure of the MD trajectory. The BRCA2 peptide is shown in cartoon representation (yellow), with residues 2403-2412 in ball-and-stick representation and residues K2404 and K2413 shown as red and blue spheres, respectively. DMC1 protomers are shown as surfaces: protomers A, B, E, and F in grey, protomer C in green, and protomer D in orange. DMC1$_C$ residues contacting BRCA2 K2404 in the MD trajectories are highlighted in pink, whereas residues contacting BRCA2 K2413 are highlighted in light blue. The movement explaining these proximities is illustrated by the white dotted arrow. **(C)** Selected contacts are monitored with time: the smallest distance between atoms of BRCA2 and DMC1 residues is plotted as a function of time in the 3 trajectories.

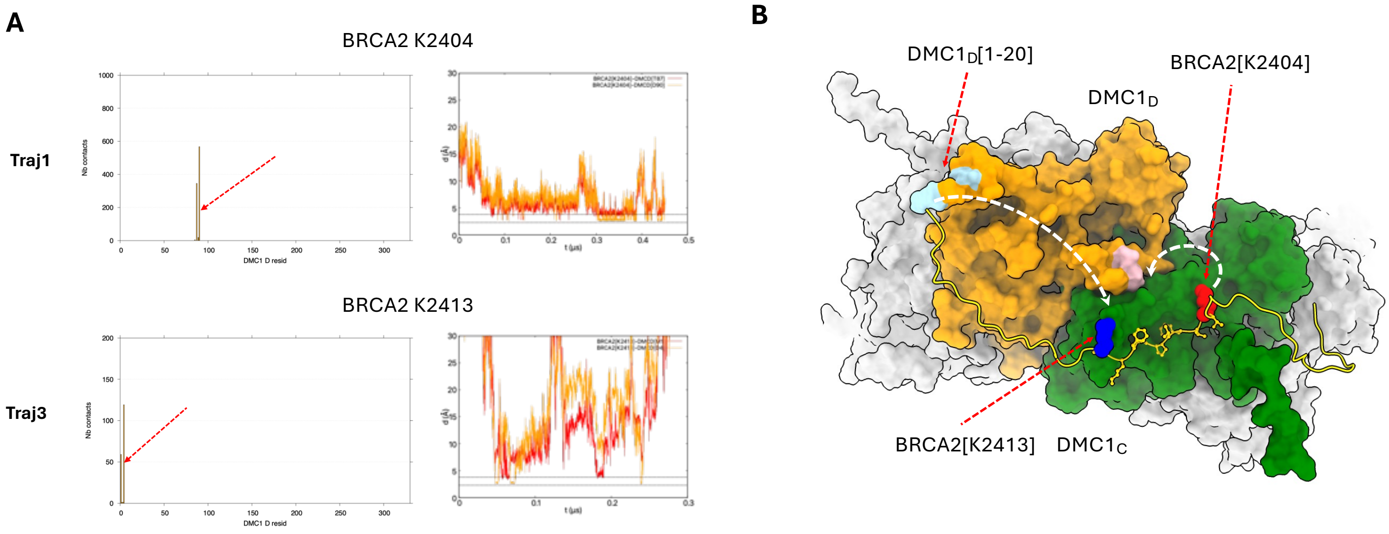


**Figure S10. Extended data on the MD simulations of Figure 5. Contact analysis of BRCA2-PL K2404 and K2413 in molecular dynamics simulations of the ssDNA–DMC1-BRCA2-PL complex: interactions with $DMC1_D$. (A)** Residues of the $DMC1_D$ protomer contacting BRCA2 residue K2404 or K2413 are shown together with the number of frames in which each contact (d<4.0Å) was detected. Frames were sampled every 0.1 ns throughout the simulations. On the right, the corresponding distance monitoring along the MD trajectories is shown for K2404 in trajectory 1 and for K2413 in trajectory 3. **(B)** Contacts are mapped onto the starting structure of the MD trajectory. The BRCA2 peptide is shown in cartoon representation (yellow), with residues 2403-2412 in ball-and-stick representation and residues K2404 and K2413 shown as red and blue spheres, respectively. DMC1 protomers are shown as surfaces: protomers A, B, E, and F in grey, protomer C in green, and protomer D in orange. $DMC1_D$ residues contacting BRCA2 K2404 in the MD trajectories are highlighted in pink, whereas residues contacting BRCA2 K2413 are highlighted in light blue. The movement explaining these proximities is illustrated by the white dotted arrow.

**Table S1**. BLI results obtained with ssDNA coated on a streptavidin biosensor.

| Nucleoproteinfilaments | BRCA2 -P | RAD54B-P |
|---|---|---|
| DMC1 – ssDNA expt 1 | **7.4E-06 ± 1.0E-6 M** | **7.3-06 ± 1.6E-6 M** |
| DMC1 – ssDNA expt 2 | | **9.2E-06 ± 2.3E-6 M** |
| DMC1 F89A – ssDNA expt1 | **7.0E-06 ± 1.0E-6 M** | **11.0E-06 ± 1.0E-6 M** |
| DMC1 F89A – ssDNA expt 2 | **7.5E-06 ± 0.4E-6 M** | **11.0E-06 ± 1.3E-6 M** |
| DMC1 D175K – ssDNA expt1 | **15.0E-06 ± 1.9E-6 M** | **21.0E-06 ± 0.8E-6 M** |
| DMC1 D175K – ssDNA expt2 | **15.0E-06 ± 1.0E-6 M** | **> 30.0E-06 M** |
| DMC1 D180K – ssDNA expt1 | **30.0E-06 ± 7.2E-6 M** | **> 30.0E-06 M** |
| DMC1 D180K – ssDNA expt2 | **> 30.0E-06 M** | **> 30.0E-06 M** |
| DMC1 E213K – ssDNA expt1 | not detected | not detected |
| DMC1 E213K – ssDNA expt2 | not detected | not detected |
| DMC1 – ssDNA expt 3 | **13.0E-06 ± 2.6E-6 M** | **14.0E-06 ± 2.9E-6 M** |
| DMC1 – ssDNA expt 4 | **9.8E-06 ± 2.0E-6 M** | **10.0E-06 ± 0.6E-6 M** |
| DMC1 Δ18 – ssDNA expt 1 | **23.0E-06 ± 2.3E-6 M** | **> 30.0E-06 M** |
| DMC1 Δ18 – ssDNA expt 2 | **25.0E-06 ± 6.3E-6 M** | **> 30.0E-06 M** |

**Table S2**. CryoEM data collection and model statistics.

| | ssDNA-DMC1 | ssDNA-DMC1-BRCA2-PL | ssDNA-DMC1-RAD54B-PL |
|---|---|---|---|
| Microscope | Titan Krios | Titan Krios | Titan Krios |
| Voltage (keV) | 300 | 300 | 300 |
| Detector | Falcon 4I | Falcon 4I | Falcon 4I |
| Magnification | 165 000 | 165 000 | 165 000 |
| Defocus range | -2.2 µM to -0.6 µM | -2.2 µM to -0.6 µM | -2.2 µM to -0.6 µM |
| Frames/movie | 30 | 30 | 30 |
| Pixel size (Å) | 0.745 | 0.726 | 0.726 |
| Electron dose (e/Å$^2$) | 50 | 50 | 50 |
| Exposure (s) | 3.48 | 2.08 | 3.84 |
| Micrographs | 8 968 | 6 884 | 10 228 |
| Picked particles | 1.9 million | 2.1 millions | 1.5 million |
| Final particles | 267 809 | 197 574 | 409 790 |
| Processing method | Helical refinement | Helical refinement | Helical refinement |
| Twist (°) | 55.6 | 55.6 | 55.7 |
| Rise (Å) | 15.9 | 15.5 | 15.9 |
| Resolution (Å) | 2.16 | 1.90 | 2.04 |
| Real-space refinement | | | |
| Composition | | | |
| Protein (residues) | 316 | 327 | 326 |
| ssDNA (nucleotides) | 3 | 3 | 3 |
| Calcium ions | 2 | 2 | 2 |
| Water molecules | 51 | 29 | 69 |
| Map model FSCaverages | 0.7811 | 0.8070 | 0.8338 |
| Bonds (rmsd) | | | |
| Lengths (Å) | 0.0065 | 0.0074 | 0.01 |
| Angles (°) | 1.671 | 1.563 | 1.891 |
| MolProbity score | 0.70 | 0.92 | 0.88 |
| Clash score | 0 | 1.12 | 0.57 |
| Rotamer outliers (%) | 0.39 | 0.36 | 0.75 |
| Ramachandran plot | | | |
| Outliers (%) | 0 | 0 | 0 |
| Favored (%) | 96.79 | 97.51 | 96.88 |
| B factors (Å$^2$) | | | |
| Median | 56.7 | 50.9 | 69.4 |
| Min | 28.4 | 25.8 | 40.3 |
| Max | 168.7 | 161.7 | 192.9 |
